# A semi-empirical model of the aerodynamics of manoeuvring insect flight

**DOI:** 10.1101/2020.01.09.900654

**Authors:** Simon M. Walker, Graham K. Taylor

## Abstract

Blade element modelling provides a quick analytical method for estimating the aerodynamic forces produced during insect flight, but such models have yet to be tested rigorously using kinematic data recorded from free-flying insects. This is largely because of the paucity of detailed free-flight kinematic data, but also because analytical limitations in existing blade element models mean that they cannot incorporate the complex three-dimensional movements of the wings and body that occur during insect flight. Here, we present a blade element model with empirically-fitted aerodynamic force coefficients that incorporates the full three-dimensional wing kinematics of manoeuvring *Eristalis* hoverflies, including torsional deformation of their wings. The two free parameters were fitted to a large free-flight dataset comprising *N* = 26, 541 wingbeats, and the fitted model captured approximately 80% of the variation in the stroke-averaged forces in the sagittal plane. We tested the robustness of the model by subsampling the data, and found little variation in the parameter estimates across subsamples comprising 10% of the flight sequences. The simplicity and generality of the model that we present is such that it can be readily applied to kinematic datasets from other insects, and also used for the study of insect flight dynamics.

## 1. Introduction

The unsteady aerodynamics of insect flight have been the focus of considerable research, with new aerodynamic mechanisms still being discovered [4]. Dipteran flies have received particular attention, because their possession of only two functional wings halves their kinematic complexity relative to four-winged insects, and avoids the aerodynamic complexity of tandem wing-wing interactions. Nevertheless, like most other insects, flies use high angles of attack and rapid wing rotation at stroke reversal, posing substantial challenges for aerodynamic modelling. Various modelling approaches have been utilised, each with their own advantages and disadvantages. More sophisticated techniques include the use of mechanical flappers [5–7] or computational fluid dynamics (CFD) [e.g. 8,16,31,42]. Both approaches determine the aerodynamic forces from a predefined set of wing kinematics, allowing the effect of specific kinematic parameters to be investigated experimentally [7,30,42], but recreating an insect’s wing kinematics in a mechanical or computational model is not straightforward, because the wings follow complex three-dimensional paths, and undergo substantial deformation through the wingbeat [36,38]. Wing deformation is difficult to replicate accurately in a mechanical flapper, and substantial user effort is required to generate a mesh capable of accommodating wing deformation in a computational model. In addition, the aerodynamics of the wings are affected by the body’s motions during flight manoeuvres, which are difficult to reproduce in a mechanical flapper and demanding to model computationally. Although the development of efficient algorithms and the increasing availability of low-cost clusters is making computational approaches ever more practical, their application is still limited to quite small datasets.

A simpler approach is to use an analytical blade element model to estimate the aerodynamic forces, by splitting the wing into a series of narrow chordwise elements, each of which is modelled independently [11]. To the extent that the flow around a real wing is inherently three-dimensional and coupled to the wake [27], a blade element model cannot capture all of the details of the unsteady flow in the way that a computational model can. Nevertheless, as we demonstrate here, it is still possible to use this approach to make practically useful predictions of the forces by using empirically-fitted force coefficients to summarise the complexities of the aerodynamics. Current blade element models comprise a quasi-steady component capturing how the pressure forces depend on the instantaneous velocity and angular velocity of the wing, and an unsteady component capturing how the pressure forces depend on the wing’s instantaneous acceleration [26,31,34], through the phenomenon of added mass. Other unsteady effects relating to the development of the flow are not captured by these models, because their coefficients are time-invariant and neglect the effects of wing-wake interactions from one half-stroke to the next [31]. Analytical blade element models therefore simplify the unsteady three-dimensional aerodynamics of flapping flight substantially, but can still do a surprisingly good job of approximating the forces produced by a flapping wing [26,31,34]. The key is to identify an appropriate analytical formulation describing how the aerodynamic forces on each blade element vary with the wing kinematics, which is the aim of this paper.

Previous studies have adopted several *ad hoc* approaches to modelling different aspects of the aerodynamics — especially the effects of wing rotation, which have been treated separately from the effects of wing translation in almost all previous models (16,26,31,34; but see 35). In each case, these models have been parameterised by a set of empirical force coefficients fitted to measurements made under different kinematic conditions using either a mechanical flapper [16,31,34] or a numerical model [26]. The resulting blade element models involve either one [16,31,34] or two [26] fitted parameters to describe the rotational lift and drag, together with separate expressions for the translational lift and drag coefficients as functions of the angle of attack, with two [26] to four [31,34] or five [16] fitted parameters. Consequently, the lift and drag are predicted from analytical expressions that are sometimes quite far removed from the underlying physics, and which involve from six [26] to nine [31,34] or even eleven [16] empirically-fitted parameters. This brings an attendant risk of over-fitting, and as the identification and verification of these models has only been done using flat, rigid wings and simplified kinematics for the simplest case of equilibrium hovering flight [16,26,31,34], it is unknown how well they predict the aerodynamic forces and moments on real insects undergoing free-flight manoeuvres involving complex wing deformations. A particular concern with using such multi-parameter models of *ad hoc* form is that these may not generalize well to other flight morphologies, wing kinematics, or flow conditions beyond those under which the data were collected. What is needed instead is a standard form developed from first principles–and therefore expected to generalize–that is capable by default of capturing the full complexity of the deforming wing kinematics.

In fact, as we show here, it is possible to predict approximately 80% of the variation in the stroke-averaged forces of free-flying hoverflies by using a physics-based model with just two free numerical parameters that are fitted empirically to the data. We achieve this by using linear least squares modelling to fit the numerical coefficients of an unsteady blade element model developed from first principles to a free-flight dataset recording the deforming wing kinematics and stroke-averaged body accelerations for *N* = 26, 541 wingbeats. Fitting this simple physics-based model to our free-flight dataset also allows us to interpret some of the more complicated empirical functions that have been fitted to model the aerodynamic force coefficients previously [6,16] and that have been co-opted into subsequent models [31,34]. Our new analytical blade element model takes full account of the three-dimensional motion of the wings and body, incorporating wing deformation in the form of a linear time-varying wing twist distribution, which is sufficient to capture most of the deformation that is present in *Eristalis* [38]. Whilst our empirical data from hoverflies do not allow us to verify how accurately our model predicts the time history of the aerodynamic forces within a single wingbeat, the stroke-averaged forces are modelled closely. Given that the body dynamics of *Eristalis* are too slow to depend closely on the periodic forcing experienced through the wingbeat [40], our model’s ability to fit the stroke-averaged forces closely makes it well-suited to use in future analyses of flight dynamics and control in this species. The simplicity and generality of our model is such that it can also be applied to kinematic datasets from other insects. Indeed, for the largest insects, in which the body dynamics operate on a similar timescale to the wingbeat [40], it should even be possible to fit the aerodynamic force coefficients directly to the time-varying rather than stroke-averaged forces.

## 2. Methods

The overall aim of this paper is to develop an aerodynamic model with empirically fitted coefficients that predicts the stroke-averaged aerodynamic forces as a function of the instantaneous kinematic state of the wing through the stroke. The empirical measurements that we analyse here are those previously described in [39], but we begin by providing a brief summary of the experimental methods and kinematic reconstruction technique for context, before providing a detailed description of the aerodynamic modelling that forms the primary contribution of this paper.

### (a) Experimental methods

Adult *Eristalis tenax* and *E. pertinax* (Diptera: Syrphidae) were caught in Oxford and released singly inside a 1m diameter opaque acrylic sphere. Four synchronised high-speed video cameras (SA3, Photron Ltd, Bucks, UK) with 180mm macro lenses (Sigma Imaging Ltd, Welwyn Garden City, UK) were used to record 768*×*640 pixel images at 3800 Hz. Bright back-illumination was provided by two synchronised 200W infrared pulsed lasers (HSI-5000, Oxford Lasers Ltd, Oxford, UK), each of which was routed through a split liquid light guide before being collimated by one of four large Fresnel lenses. The 805 nm wavelength of the laser was far beyond the range of the visible spectrum for *Eristalis* [upper limit: 600 nm; 1,2,17], and a 20 *µ*s pulse duration was used to eliminate motion blur and to prevent overheating of the insect. Ambient lighting was provided by an overhead LED light source. The cameras were self-calibrated using custom-written software in MATLAB (The Mathworks Inc., Natick, MA, USA), to identify jointly optimal estimates of the camera parameters and the spatial coordinates of points on a 2D calibration grid held in a range of positions and orientations [37]. We captured between 10 and 50 separate recordings of each hoverfly, before anaesthetising it with CO_2_ at the end of the experiment, and weighing it using a microbalance with 0.1*µ*g readability (UMX2, Mettler Toledo Ltd, Leicester, UK). We analysed all of the flight sequences in which both wings were visible to all four cameras for *≥* 5 wingbeats, giving a total sample of *N* = 26, 541 wingbeat pairs from 854 flight sequences representing 36 hoverflies. A fully automated shape-carving procedure was used to label voxels contained within the minimum convex hulls of the insect’s body and its two wing outlines, respectively [39].

### (b) Flight kinematics modelling

Throughout the paper, we use boldface symbols to represent vector quantities, and use *t* to represent continuous time. We defined the body kinematics using a right-handed, rotating, body-fixed axis system *{x*_*b*_, *y*_*b*_, *z*_*b*_*}* with its origin at the centre of volume of the body voxels, its *x*_*b*_-axis directed anteriorly along the major axis of the body voxels, and its *y*_*b*_-axis pointing rightward parallel to the line connecting the wing bases (Fig. 1). We measured the position of the body axes, ***X***_*b*_(*t*), in a non-rotating laboratory coordinate system *{X, Y, Z}*, in which the *Z*-axis was vertical, and smoothed our measurements using a quintic spline fitted in B-form in Matlab [29]. This method fits each element of the position vector as the smoothest piecewise polynomial function of time that falls within a given tolerance of the data, defined as the sum of the squared distance of the function from the data over all sample points. For transparency, we selected a tolerance that was equal to the sum of the squared variation that would have been removed by forward-backward filtering the data using a 3rd order Butterworth filter with a 100 Hz cut-off frequency (−3 dB) chosen to fall well below the insect’s 188 ± 14 Hz wingbeat frequency (mean ± SD). We then double-differentiated each quintic spline analytically in Matlab to estimate the insect’s instantaneous linear acceleration with respect to the laboratory coordinate system, 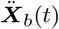, resolved in the lab axes.

**Figure 1.**
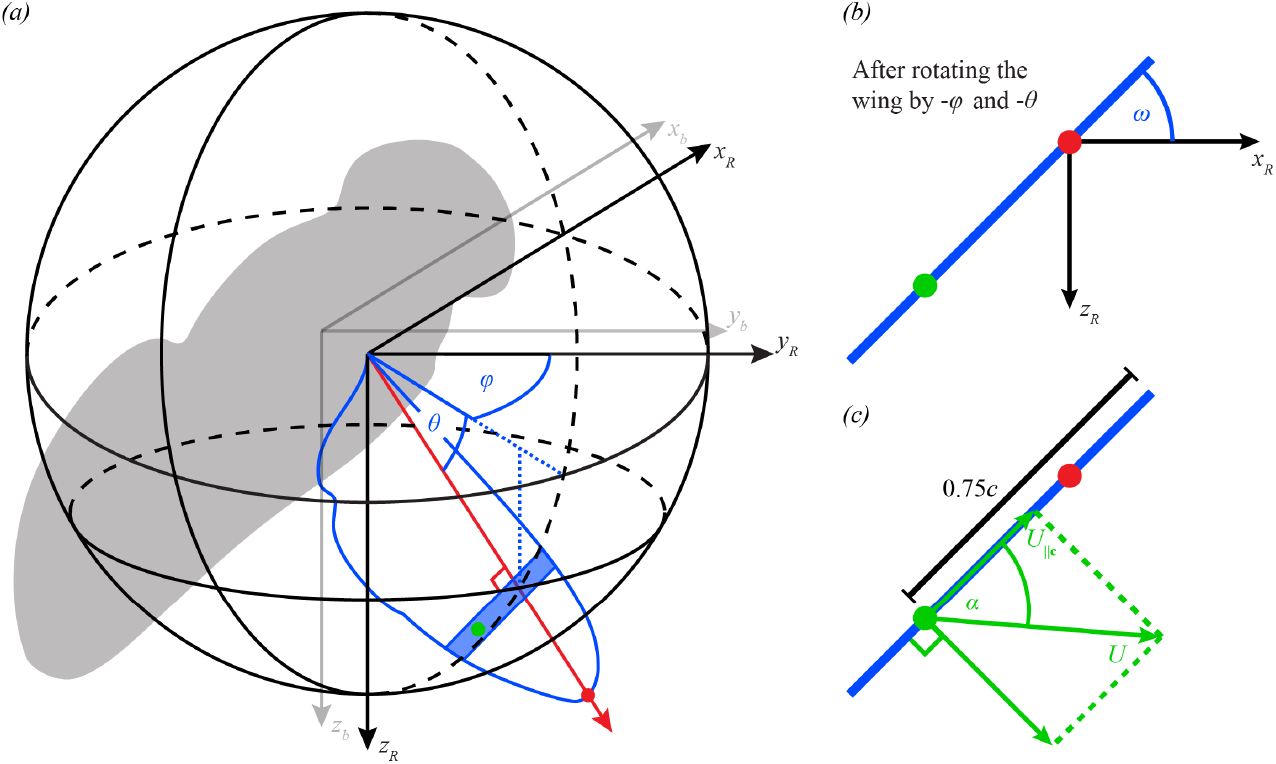
Kinematic definitions. *(a)* The body axes *{x*_*b*_, *y*_*b*_, *z*_*b*_*}* are defined as a right-handed axis system (grey arrows) with its origin at the centre of volume of the insect’s body (grey silhouette), its *x*_*b*_-axis directed anteriorly along the major axis of the body voxels, and its *y*_*b*_-axis pointing to the insect’s right, parallel to the line connecting the wing bases. The tip kinematics of the right wing are defined by its spherical coordinates in another right-handed axis system *{x*_*R*_, *y*_*R*_, *z*_*R*_*}* (black arrows) fixed parallel to the body axes, but originating at the wing base. The tip kinematics of the left wing are defined by its spherical coordinates in an equivalent left-handed axis system *{x*_*L*_, *y*_*L*_, *z*_*L*_*}* (not shown). In each case, the line from wing base to wing tip (red dot) defines the spanwise rotational axis of the wing (red arrow): the azimuth of this spanwise axis defines the stroke angle of the wing (*ϕ*), and its elevation defines the deviation angle (*θ*). The blue shaded area shows a single chordwise blade element with the position its three-quarter chord point marked by a green dot. *(b)* The pitch angle *ω* of a blade element is defined having first rotated the wing’s measured outline through its stroke angle −*ϕ* about the *z*_*R*_-axis, then through its deviation angle −*θ* about the *x*_*R*_-axis, so as to bring the line from wing base to wing tip into alignment with the *y*_*R*_-axis. The pitch angle *ω* was then defined as the angle from the *x*_*R*_*y*_*R*_-plane to the anatomical ventral side of the rotated chord perpendicular to the *y*_*R*_-axis. *(c)* The speed of a blade element (*U*) is measured at its three-quarter chord point (green dot), and is defined as the hypotenuse of the components of the velocity vector directed parallel to the blade element chord (*U*_‖*c*_) and normal to the blade element surface (*U*_⊥*S*_). The aerodynamic angle of attack (*α*) is defined as shown by the arctangent of these two vector components.

We described the kinematics of the right wing in a right-handed axis system *{x*_*R*_, *y*_*R*_, *z*_*R*_*}* parallel to the rotating body axes *{x*_*b*_, *y*_*b*_, *z*_*b*_*}*, but with its origin at the wing base. We used the spherical coordinates of the wing tip to define the stroke angle (*ϕ*) and deviation angle (*θ*) of the wing (Fig. 1). We defined the local pitch angle *ω*(*r*) of the wing at radial coordinate *r* (Fig. 2) by rotating the wing’s measured outline through the angle ϕ about *z*_*R*_, then through the angle −*θ* about *x*_*R*_, so as to bring its spanwise axis into alignment with *y*_*R*_. The local pitch angle *ω*(*r*) of the wing was then defined as the angle from the *x*_*R*_*y*_*R*_-plane to the anatomical ventral side of the chord, perpendicular to the *y*_*R*_-axis at radial coordinate *r* (Figs. 1B, 2). We summarised the spanwise twist as a linear function of distance along the wing by regressing *ω*(*r*) on *r*, modelling the local pitch angle as:

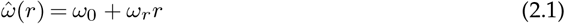

where *ω*_0_ is the pitch angle offset, and *ωr* is the linear twist gradient. The kinematics of the left wing were defined independently using an equivalent left-handed axis system *{x*_*L*_, *y*_*L*_, *z*_*L*_*}*. Variable wing camber might also have been present [38], but measuring this requires data of higher order than can be obtained using a voxel carving method to identify the wing outlines. Were wing camber to be measured directly in a future study, it would in principle be possible to incorporate its effects in the blade element model below by treating camber as augmenting the aerodynamic angle of attack defined with respect to the angle of the chord line measured between the leading and trailing edge.

**Figure 2.**
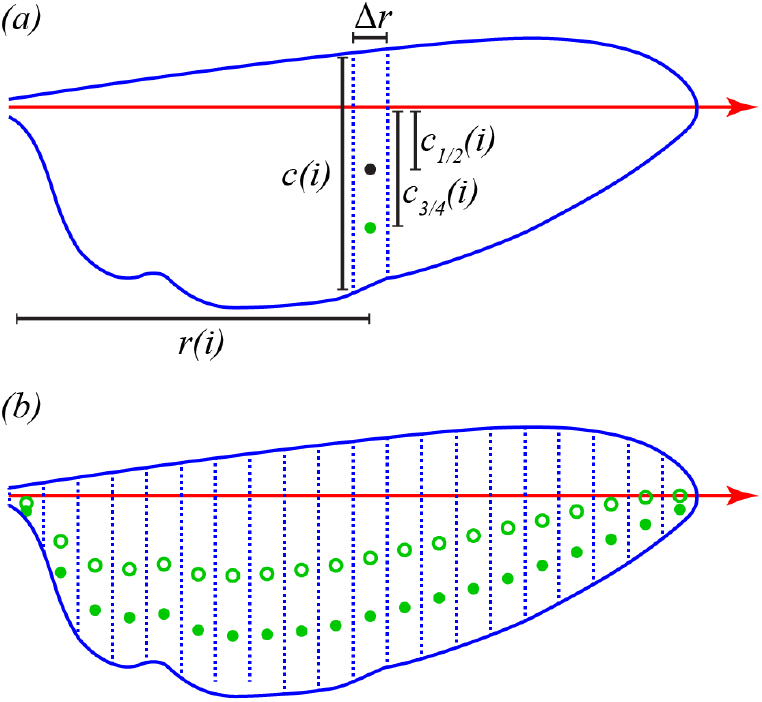
Morphological definitions and measurements. *(a)* The wing is split into twenty equally-spaced chordwise blade elements, each of width Δ*r*. The *i*th blade element is located at radial distance *r*(*i*) from the wing base, and has chord length *c*(*i*). The distance of the three-quarter chord point (open circle) from the spanwise rotation axis of the wing (black arrow) is denoted *c*_3*/*4_(*i*), and is used in calculating the quasi-steady lift and drag forces (see Supplementary Methods). Likewise, the distance of the half-chord point (closed circle) from the spanwise rotation axis is denoted *c*_1*/*2_(*i*), and is used in calculating the added mass force (see Supplementary Methods). *(b)* The measured positions of the three-quarter chord points (open circles) and half-chord points (filled circles) are shown for each of the twenty blade elements.

We smoothed our measurements of *ϕ, ω*_*r*_, and *ω*_0_ for each wing using the same quintic spline method as we used to smooth the body kinematics, this time setting the tolerance to give a similar degree of smoothing to a digital Butterworth filter with a cutoff frequency of 500 Hz for the wing tip kinematics *θ* and *ϕ* and 800 Hz for the wing twist kinematics *ωr* and *ω*_0_ (Fig. S1). We then evaluated the first and second derivatives of these fitted spline functions analytically. The smoothed kinematic data were next split into discrete wingbeats by identifying the time at which the mean angular speed of the two wing tips reached a minimum at the end of each half stroke. Finally, for consistency of sampling between discrete wingbeats of variable period, we used cubic interpolation to resample the smoothed wing and body kinematic data and their time derivatives at 100 evenly-spaced time steps through each wingbeat, beginning at the start of the downstroke. This means that the kinematic measurements were upsampled by approximately a factor of 5 prior to further analysis, which: (i) ensures that each wingbeat starts and ends at exactly the same phase; (ii) allows the data to be stored in an efficient matrix form; and (iii) standardises the basis on which the wingbeat averaged forces are estimated.

By the end of this process, each wingbeat is represented by 4,200 datapoints, comprising the 6 degrees of freedom of the body and 4 primary kinematic variables of each wing, each sampled together with their first and second derivatives 100 times per wingbeat. In principle, these data are already in a form suitable for the subsequent blade element analysis. However, in the interests of compressing the data into a compact functional form suitable for sharing, and noting the different characteristic timescales on which the different kinematic components vary, we projected the data for each wingbeat into a set of harmonic basis functions [25] comprising a truncated Fourier series plus cubic polynomial in time from the start of the wingbeat, with harmonic content to 1^st^ order for the body kinematics, 4^th^ order for the wing tip kinematics, and 6^th^ order for the wing twist kinematics (see Supplementary Methods). This compression reduces the dimension of the data by almost a factor of 30, whilst preserving *>* 99.99% of the measured variation in the pose of the insect, as characterised by the 6 degrees of freedom of its body and 4 primary kinematic variables of each of its wings. These harmonic representations of the data are shared as Supporting Data S1, so to ensure the repeatability of our analysis and to enable its validation using numerical techniques possibly requiring finer time steps, we obtained the 4,200 datapoints that we use in the blade element model for each wingbeat by evaluating these harmonic fits rather than the spline fits on which they are modelled. This step makes a negligible difference to the numerical values of the datapoints used in the analysis (Fig. S1), and hence to the results of the analysis, but it aligns the present work more closely to the approaches that we have developed elsewhere for analysing the dominant kinematic couplings involved in insect flight control using harmonic functional principal components analysis [25].

### (c) Standard hovering wingbeat

We defined a standard wingbeat for use in model validation by taking the mean through time of the 1% of all *N* = 26, 541 wingbeat pairs that met most closely the criteria for hovering flight. We selected these as the 265 wingbeats with the lowest flight speed from within the set of wingbeats representing near-equilibrium flight, which we defined as flight where the magnitude of the body’s acceleration was < 0.5ms^−2^ in both the vertical and the horizontal. (i.e. such that the vertical aerodynamic force would have been within 5% of exactly supporting body weight). The 265 wingbeats that we used in this averaging came from 51 different flight recordings and 19 different individuals, and the standard hovering wingbeat that they define should therefore be representative of equilibrium hovering flight (Fig. 5).

### (d) Flight dynamics modelling

The overall goal of this paper is to use aerodynamic modelling to relate free-flight measurements of body kinematics to the wing kinematics that produce them. Although we take full account of the body’s rotational and translational motion in defining the motion of the wings relative to the air, our modelling of the resulting aerodynamic forces only considers their effects on the translational motion of the centre of mass. That is to say, we do not attempt to model the rotational dynamics of the body, which is a more complex problem requiring knowledge of the insect’s inertia tensor and the chordwise position of the centre of pressure. Subject to making such further assumptions, the rotational dynamics can be modelled subsequently using the same blade element model if required.

The only ways in which a fluid can impart force to a solid surface are through pressure forces acting normal to the surface, and friction forces acting tangential to it. We may therefore use Newton’s Second Law to write the equations of translational motion for a free-flying insect as:

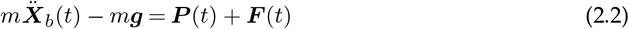

where *m* is the insect’s mass, 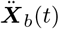 is the acceleration of the insect’s centre of mass with respect to the laboratory coordinate system, ***g*** is gravitational acceleration, ***P*** is the total pressure force, and ***F*** is the total friction force. All of these vector quantities are resolved in the non-rotating axes of the lab-fixed coordinate system, and they take full account of the angular velocity of the centre of mass and of the centripetal acceleration that produces this during manoeuvres. It is important to note that Eq. 2.2 properly contains only the external forces acting at the insect’s centre of mass, so does not show the inertial forces that act in reaction to flapping at the wing hinge. These are internal forces which cannot therefore produce any acceleration of the insect’s centre of mass, so this is a very different situation to that which is encountered when measuring the internal forces on a bench-mounted mechanical flapper. Moreover, although the wings’ motion causes variation

in the anatomical position of the centre of mass that can cause the body to oscillate at wingbeat frequency in some large insects [41], the wings are several orders of magnitude lighter than the body in *Eristalis* and therefore have a negligible effect on our measurement of the motion of the insect’s centre of mass by tracking the body.

Since the wingbeat period of a hoverfly is much shorter than any characteristic timescale of its body’s dynamics [40], we may reasonably model its body dynamics using the stroke-averaged versions of these variables, which we denote using overbar notation with *n* as wingbeat number:

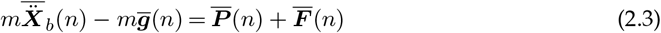

This averaging is beneficial in removing the noise associated with estimating the double derivative of position from high-speed videogrammetric data. Nevertheless, in future work with larger insects whose body dynamics operate on a similar timescale to the wingbeat [40], it might be more appropriate to retain the original form of Eq. 2.2. The lefthand side of Eq. 2.3 is a direct empirical estimate of the stroke-averaged aerodynamic forces, which we refer to hereon as the “measured” aerodynamic force, to distinguish this from the indirect estimates of the time-varying aerodynamic force that we make later using our blade element model. The stroke-averaged expression in Eq. 2.3 forms the basis of the empirical estimation of the aerodynamic forces in this paper.

### (e) Aerodynamic modelling

We begin this section by analysing the scaling of the aerodynamic forces, which we use to define the theoretical form of the kinematic predictor variables that we use to fit the aerodynamic coefficients of the quasi-steady blade element model. Although the key concepts are covered in aerodynamics texts such as Katz & Plotkin [19] and in the seminal work on insect flight by Ellington [11], they are usually only discussed in the context of detailed aerodynamic models that aim to determine analytically the same force coefficients as we aim to estimate empirically, under restrictive assumptions that will not be satisfied in insect flight. Hence, following the approach advocated by [33], and noting that there is no analytical theory which covers the full range of flow conditions experienced in insect flight, we here develop the scaling of the aerodynamic forces from first principles, with the goal of making the assumptions of our blade element model fully transparent. We then introduce the theoretical form of the equations that we use to model the unsteady forces, before describing how we fit the empirical aerodynamic force coefficients to the measured flight data. Although some elements of our model are shared with previous work, this approach results in an aerodynamic model with a different mathematical form to the others that have been used previously to analyse insect flight [6,11,16,26,31,34].

### (i) Functional form of the quasi-steady aerodynamic forces

The pressure force in Eqs. 2.2–2.3 will dominate the friction force at the Reynolds numbers of order 10^3^ that characterise *Eristalis*. This implies that the shear layer surrounding the wing must be thin in relation to its chord, which in turn means that viscous shear in this inner region of the flow cannot be directly responsible for setting fluid into motion in the outer region of the flow. This fluid motion must instead be driven by the pressure gradients that result from the constraint that fluid cannot penetrate the wing’s surface. This boundary condition means that at every point on the wing’s surface, the surface-normal component of the flow (*Q*_⊥*S*_) must be the same as the surface-normal component of the wing’s own velocity (*V*_⊥*S*_), which we may write locally as:

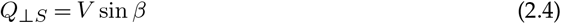

where *β* is the local angle of attack measured between the plane tangent to the surface of the wing at a given point and the inertial velocity of the wing’s surface at that point, and where *V* is the local inertial speed of the wing. The pressure gradients resulting from the global imposition of this boundary condition will modify the tangential flow at the wing’s surface, resulting in a net circulation (Γ) defined as the line-integral of the tangential flow around the airfoil. The strength of the resulting circulation cannot be determined without making further and more restrictive assumptions, but must depend on the magnitude of the normal flow in Eq. 2.4.

For a thin flat plate airfoil undergoing pure translational motion through still air, the local angle of attack *β* is the same everywhere, so is equal to the average angle of attack *α* between the velocity of the airfoil and its chord. If the airfoil is also rotating steadily, then *β* will vary linearly with distance from the axis of rotation, and we must therefore specify how to weight these local contributions to the average angle of attack *α*. For a linear weighting distribution, this is equivalent to specifying a unique point on the airfoil at which the boundary condition in Eq. 2.4 is to be satisfied, which falls three-quarters of the distance back from the leading-edge under both classical lifting-line theory and the inviscid theory of small-amplitude unsteady motion for a pitching and plunging flat plate [19]. Hence, for a flat plate airfoil undergoing quite general motions in a potential flow, the quasi-steady effects of wing rotation can be captured by defining the average angle of attack *α* as being equal to the local angle of attack *β* measured at the three-quarter chord point [11]. We take this as our starting point here, but also test this assumption empirically.

The form of Eq. 2.4 immediately suggests that under steady or quasi-steady conditions, the circulation Γ around an airfoil can be expected to vary as a function of *cU* sin *α*, where *c* is the chord length, and where the average speed *U* and average angle of attack *α* of the airfoil are measured as the local values of *V* and *β* at the three-quarter chord point. Note that this sinusoidal dependence on *α* depends simply on the averaging of the boundary condition expressed in Eq. 2.4, and not on any more detailed assumptions. In fact, the Kutta-Joukowski theorem for two-dimensional inviscid flows predicts that the pressure force varies as *ρU*Γ where *ρ* is the fluid density [19], so this inference is also consistent with the classical result that the quasi-steady pressure force is fixed at a value *πρU*^2^ sin *α* per unit span for a two-dimensional flat-plate airfoil operating at a low angle of attack with the flow departing smoothly from the trailing edge [19]. Although there is no necessary reason to expect linearity in sin *α* under other flow conditions, any aerodynamic model that we fit should naturally be able to handle this potential flow regime in addition to the separated flow conditions applicable to insect flight. This being so, it is reasonable to propose modelling the quasi-steady pressure force (*P*_*qs*_) on a three-dimensional blade element of width Δ*r* using the scaling:

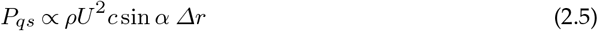

where the unknown constant of proportionality and direction of action of the force remain to be determined empirically.

The Kutta-Joukowski theorem predicts that this quasi-steady pressure force acts perpendicular to the relative velocity of a two-dimensional airfoil at low angles of attack, but some departure from this is inevitable with the separated flows experienced on three-dimensional wings at higher angles of attack. In fact, the pressure force acts more nearly perpendicular to the chord when the flow is separated over the upper surface of the wing [19]. This contradiction is not too problematic in practice, because even at low angles of attack, the flow induced by the wake tilts the aerodynamic force vector back considerably on a low aspect ratio wing, bringing its line of action more nearly perpendicular to the chord. In any case, given that the main contributions to the stroke-averaged pressure force come when the wing is at very high angles of attack, it is reasonable to assume—as a first approximation—that the quasi-steady pressure force acts perpendicular to the chord in the direction of increasing angle of attack. Decomposing this force into a quasi-steady lift component, *L* = *P*_*qs*_ cos *α*, and a quasi-steady drag component, *D* = *P*_*qs*_ sin *α*, then yields the scaling:

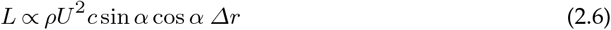

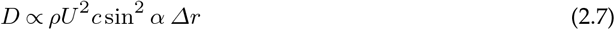

where the constant of proportionality is assumed to be the same in both expressions, and where by definition the lift acts perpendicular to the relative flow and the drag in the direction of it.

### (ii) Quasi-steady force coefficients

By convention, the non-dimensional lift coefficient (*C*_*L*_) and drag coefficient (*C*_*D*_) of a blade element are defined as:

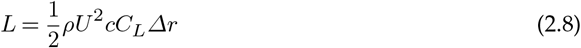

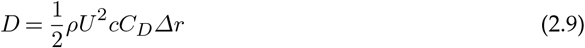

and similarly for the pressure force coefficient (*C*_*P*_) with respect to the overall quasi-steady pressure force *P*_*qs*_. Combining these identities with the scalings in Eqs. 2.6–2.7 implies that the local lift and drag coefficients may be approximated as:

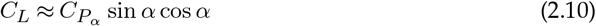

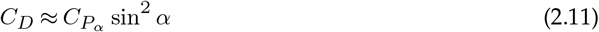

where the unknown constant 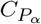 remains to be determined empirically, but can be interpreted as the derivative of the pressure force coefficient *C*_*P*_ with respect to the angle of attack *α*, at vanishing angle of attack.

These approximations imply that lift and drag should both vanish at *α* = 0, which reflects the neglect of friction to this point. However, whereas the pressure force is expected to dominate the friction force at most angles of attack, friction cannot be dismissed entirely at very low angles of attack, for which the pressure force will be small and the friction force tangent to the wing’s surface may be the primary contributor to drag. Because friction depends only on the tangential flow within the boundary layer, it does not depend strongly on the angle of attack, except at very high angles of attack when its contributions are in any case negligible. Hence, whilst the component of the tangential force resolved in the direction of drag will in principle vary with cos *α*, we may make use of the fact that cos *α* ≈ 1 at low angles of attack and approximate the effects of friction by adding a small constant offset to the righthand side of Eq. 2.11, such that:

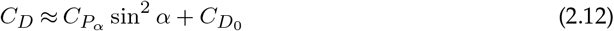

where 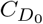 denotes the value of the drag coefficient at zero angle of attack. This approach neglects possible variation of the friction drag coefficient with Reynolds number through the stroke, but this is unlikely to represent much of a limitation in practice, given that the offset term 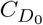 is determined empirically, and lumps the friction drag together with any pressure drag that may also happen to be present at zero angle of attack owing to the effects of wing corrugations, leading-edge thickness, etc. Such effects are difficult to model from first principles, being highly dependent on the detailed wing structure.

### (iii) Functional form of the unsteady forces

The quasi-steady aerodynamic force must be supplemented by an unsteady aerodynamic force whenever the wing accelerates, which can be shown by rewriting the normal component of the local flow velocity in Eq. 2.4 as *V*_⊥*S*_ = *V* sin *β*, and differentiating this restated boundary condition with respect to time *t* to yield:

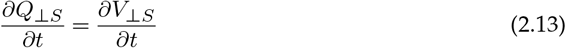

which shows that any acceleration of the wing in the direction normal to its surface must be accompanied by an equal acceleration of the fluid at the wing’s surface. In accordance with Newton’s laws of motion, the reaction to this acceleration of the fluid must be an unsteady pressure force (*P*_*us*_) acting in the opposite direction normal to the wing’s surface. Naturally, the surrounding fluid must also be accelerated, to a degree that will decline away from the wing, but the strength of the reaction force can be expected to scale linearly with the normal component of the wing’s acceleration, so that the resulting forces will vary as if the wing were accelerating a virtual mass of fluid with it. For this reason, the unsteady pressure force is usually referred to as an added mass force.

The added mass force is complicated to model for a body of arbitrary shape undergoing arbitrary motion, and the added mass on insect wings has conventionally been approximated using the potential flow solution for a flat plate in translational motion at a constant rate of acceleration [11,26,31,34]. This gives the the unsteady pressure force for any given blade element as:

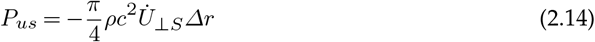

where 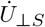 is the acceleration of the blade element normal to its surface, defined as the projection of its acceleration onto the surface normal, signed positive in the direction of increasing angle of attack. Note that whereas the velocity of a blade element is defined at the three-quarter chord point for the purposes of determining the quasi-steady aerodynamic forces, the acceleration of the blade element is defined at the half-chord point for the purposes of determining the unsteady added mass forces. The added mass force will not sum to zero over a periodic flapping cycle, except under some specific and unrealistic symmetry conditions. Nevertheless, the stroke-averaged added mass force is expected to be small in comparison to the scale of its variation through the wingbeat, and small in comparison to the stroke-averaged quasi-steady aerodynamic force. On grounds of low signal to noise ratio, it therefore makes sense to model it directly using Eq. 2.14, rather than to attempt to model it empirically using the regression technique that we use for the quasi-steady aerodynamic forces in the next section.

### (iv) Blade element modelling

The final step in the aerodynamic modelling is to use the equations in Section 2ii to assemble a set of kinematic predictors for the stroke-averaged quasi-steady aerodynamic forces. Given our measurements of the wing and body kinematics, these results can then be used together with Eqs. 2.2 and 2.14 to formulate a set of linear equations that can be solved in a least squares sense for the unknown aerodynamic force coefficients 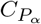 and 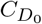, which are assumed to be the same for all blade elements. Practically speaking, we split each wing into 20 evenly-spaced blade elements, of width Δ*r* and chord length *c*(*i*), where *i* ∈ 1 … 20 denotes the blade element number (Fig. 2), the aerodynamic contributions of which we then summed over all 20 blade elements and all 100 sample points for both wings.

Making use of the equations in Section 2ii, we modelled the instantaneous lift (***L***), drag (***D***), and added mass (***A***) forces acting on the *i*th blade element at time *t* as:

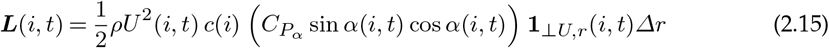

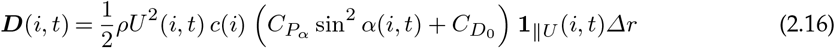

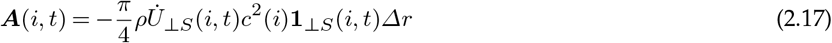

in which **1**_‖*U*_, **1**_⊥*U,r*_, **1**_⊥*S*_ are unit vectors resolved in the body axes and defined as follows. The unit vector **1**_⊥*U,r*_ is directed mutually perpendicular to the velocity of the blade element and its span, and points in the direction of increasing angle of attack, thereby defining the direction in which lift acts. The unit vector **1** ‖ _*IU*_ is directed parallel to the velocity of the blade element, and points in the direction of the relative flow, thereby defining the direction in which drag acts. In each case, the blade-element velocity is defined at the three-quarter chord point. The unit vector **1**_⊥*S*_ is the blade-element surface normal, signed positive in the direction of increasing angle of attack, and is related to the other two unit vectors by the identity **1**_⊥*S*_ = **1**_⊥*U,r*_ cos *α* + **1**_*IU*_ sin *α*. The aerodynamic speed, *U*, and angle of attack, *α*, of each blade element are each measured at the three-quarter chord point working back from the leading edge, as detailed in the Supplementary Methods, such that Eqs. 2.15 and 2.16 account implicitly for the effect of wing rotation on the quasi-steady pressure force (see also Section 2i). In contrast, the normal acceleration 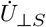 is defined at the half-chord point for the purposes of determining the added mass force (see Supplementary Methods). All of these quantities measure the blade element kinematics with respect to the laboratory coordinate system, and therefore incorporate all of the effects of the body’s translational and rotational motion during forward flight and manoeuvring.

Our kinematic measurements record the motion of each wing at *τ* ∈ 1 … 100 discrete time points through each wingbeat. The contribution of each wing to the total stroke-averaged aerodynamic force may therefore be written as:

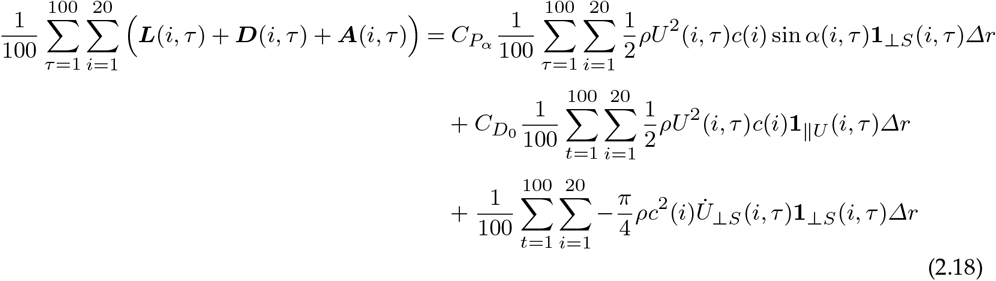

where we have made use of the identity **1**_⊥*S*_ = **1**_⊥*U,r*_ cos *α* + **1** ‖ _*IU*_ sin *α* to eliminate two of the trigonometric terms when combining Eqs. 2.15 and 2.16. By an obvious use of notation for the summations, we will abbreviate the righthand side of Eq. 2.18 as 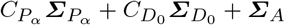. The total stroke-averaged aerodynamic force on a given wingbeat may therefore be modelled as:

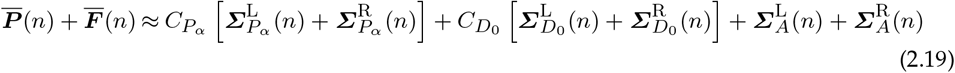

where the L and R superscripts denote summations for the left and right wings, respectively.

### (f) Model fitting

Combining Eq. 2.19 with Eq. 2.3 for the measured stroke-averaged aerodynamic force, we may write:

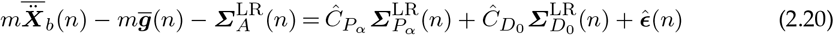

where 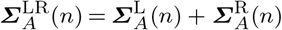 and so on, and where 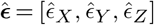 is a residual error term accounting for the difference between the measurements and the model in the lab axes in which the forces are resolved. Eq. 2.20 is linear in the unknown force coefficients 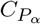 and 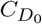, so can be solved using linear regression to provide parameter estimates 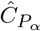and 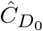 that minimise the error sum of squares 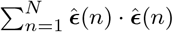 over all *N* = 26, 541 wingbeats. We initially solved Eq. 2.20 with the aerodynamic speed *U* and angle of attack *α* defined at the three-quarter chord point as explained above, which yields a unique solution for 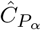 and 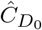. As a direct check on the effect of this assumption of the model, we then tried varying the chordwise position at which *U* and *α* were defined, which yields a family of solutions for 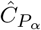 and 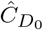. Finally, we verified the importance of including body motion and wing twist in the model by re-computing the kinematics without accounting for body motion, and with the wing pitch angle set uniformly at its value mid-span, before solving again for the unknown force coefficients 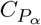 and 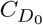 (see Table 1).

**Table 1.** Comparison of the full aerodynamic model with several alternative models, showing the effect of various simplifications on the estimated force coefficients 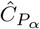 and 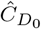, and on the mean squared error (MSE) in the predicted forces. The penultimate column reports the chordwise reference point at which the kinematics are defined in the blade element model. This chordwise reference point is optimised in the version of the full model shown in the last row of the table, so as to minimise the MSE subject to the necessary physical constraint that 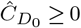. The last column gives the total number of free parameters estimated by the optimisation procedure. The MSE is averaged over all *N* wingbeats and all three axes such that 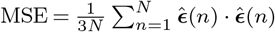, where 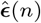 contains the residual error in each axis for the *n*^th^ wingbeat. The estimated force coefficients are reported ±1SD, where SD is the standard deviation of their values computed over 10^5^ random subsamples of the data, each comprising 10% of the recorded flight sequences.

| Model | $\hat{C}_{P_\alpha}$ | $\hat{C}_{D_0}$ | MSE (mN)<sup>2</sup> | % chord | parameters |
| --- | --- | --- | --- | --- | --- |
| full model | $2.70 \pm 0.04$ | $0.15 \pm 0.07$ | 0.0265 | 75 | 2 |
| no drag offset | $2.73 \pm 0.04$ | – | 0.0267 | 75 | 1 |
| flat plate wing | $1.69 \pm 0.04$ | $0.48 \pm 0.06$ | 0.0344 | 75 | 2 |
| no body motion | $2.65 \pm 0.04$ | $0.86 \pm 0.19$ | 0.0356 | 75 | 2 |
| % chord optimised | $2.58 \pm 0.04$ | 0 | 0.0262 | 87 | 3 |

### (g) Model validation

In light of the very large sample, and because the regression model does not take account of autocorrelation in the stroke-averaged forces from one wingbeat to the next, we do not report 95% confidence intervals for our parameter estimates. Instead, we tested the robustness of the analysis using subsampling, repeating the regression modelling 10^5^ times on random subsamples of the data each containing only 10% of the flight sequences (i.e. 85 out of the 854 flight recorded sequences). This subsampling analysis allows us to assess the variance in our parameter estimates resulting from variation between individuals and flight sequences, and does so at a sample size that is more realistic for future studies than the very large sample used here (i.e. order 10^2^ rather than order 10^3^ flight sequences). Finally, using our estimates of 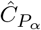 and 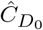 with the kinematics defined at the three-quarter chord point, we tested how the number of blade elements and number of time steps affected the aerodynamic forces predicted for the standard hovering wingbeat. The predicted mean absolute aerodynamic force changed by only 0.30% when increasing the number of blade elements to 1,000 (Fig. S2a,b), and by only 0.14% when increasing the number of time steps to 10,000 (Fig. S2c,d), so we conclude that the default discretisation using 20 blade elements and 100 time steps is more than sufficient.

## 3. Results

### (a) Body dynamics

Our free-flight dataset captures a wide variety of behaviours, including forward flight, hovering, ascent, descent, and saccadic manoeuvres. A typical flight recording (Supplementary Video S1) includes brief periods of slow forward flight, punctuated by fast body saccades. Although we cannot claim to have captured the entire flight envelope, these data therefore cover a large part of the behavioural repertoire of *Eristalis*, including many manoeuvres typical of free-flight [13,14,20,21]. This range of behaviour is reflected in the variability of the measured aerodynamic forces acting along the *x*_*b*_-axis (0.97 ± 0.43 mN) and *z*_*b*_-axis (−1.11 ± 0.4 mN) of the body (mean ± SD; Fig. 3a,c). The forces measured along the *y*_*b*_-axis were much less variable (0.00 ± 0.11 mN; Fig. 3b), consistent with the orthodoxy that comparatively little lateral aerodynamic force is produced during manoeuvres [3]. Nevertheless, the double derivatives of body position vary to a similar extent in all three body axes (Fig. 3d-f), indicating that the asymmetry of force production in the *x*_*b*_- and *z*_*b*_-axes is explained by the need to overcome the body’s acceleration due to gravity, which is small in the *y*_*b*_-axis except during highly-banked turns. It follows that after addressing the requirement for weight support, the hoverflies were actually comparably manoeuvrable in all three body axes.

**Figure 3.**
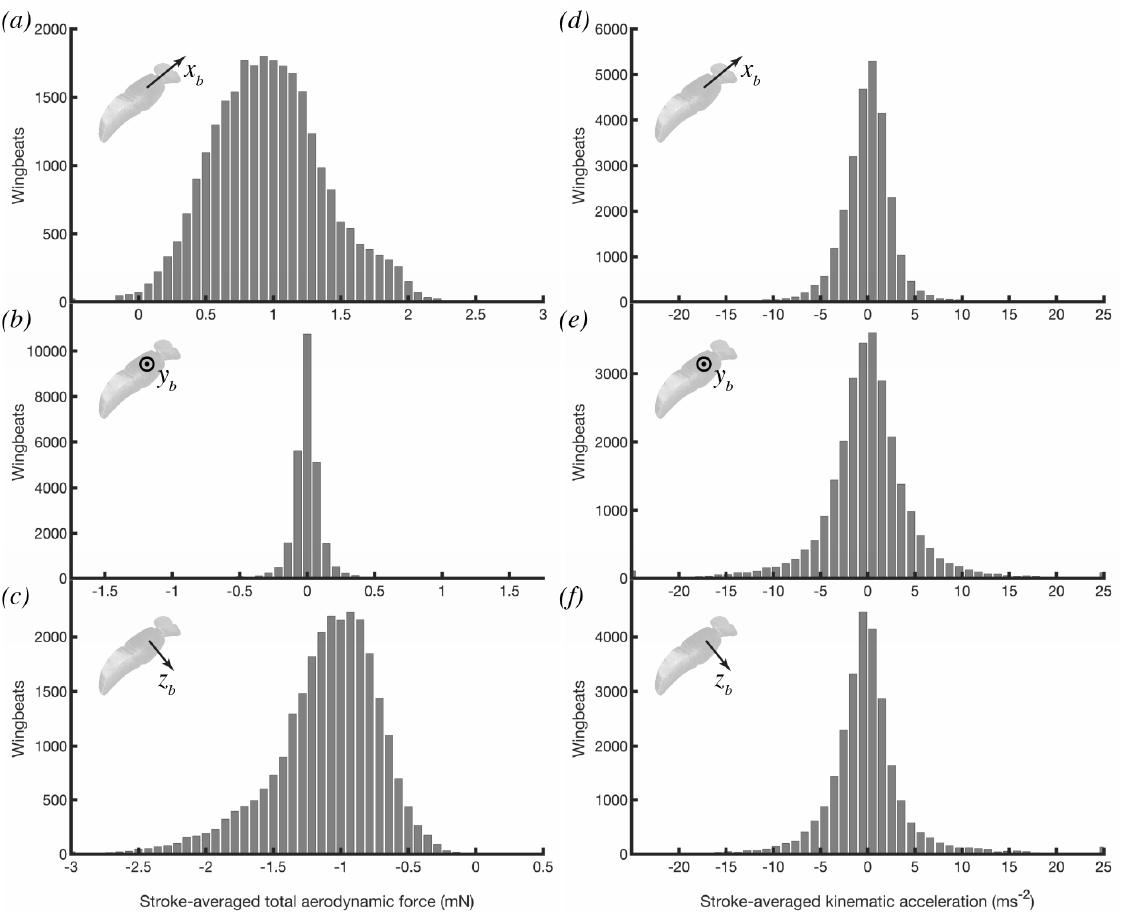
Histograms of the measured stroke-averaged aerodynamic force and measured acceleration of the insect’s body in an inertial frame of reference. (*a-c*) Measured stroke-averaged aerodynamic force, resolved in the insect’s body axes. (*d-f*) Measured stroke-averaged acceleration, 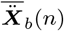, resolved in the insect’s body axes. Note that the distribution of the acceleration is similar across all three body axes, but that the need to provide weight support in addition to manoeuvring force means that the resultant forces are principally distributed in the *x*_*b*_- and *z*_*b*_-axes of the body.

### (b) Wing kinematics

Wingbeat frequency and stroke amplitude vary greatly over the dataset, but their variability owes more to variation between individuals than within (Fig. 4), and the time-history of the wing kinematics is actually quite stereotyped over the whole dataset (Fig. 5B). Because the wing tip trajectory is inclined at approximately 45°to the body, the stroke angle *ϕ* and deviation angle *θ* always have similar oscillation amplitudes, and both vary approximately sinusoidally through the wingbeat (Fig. 5A). The wing pitch angle *ω* at mid-span varies symmetrically on the upstroke and downstroke, showing a slight recoil at the start of each half-stroke. The aerodynamic angle of attack *α* has a similar time history on both the upstroke and the downstroke, changing rapidly as the wing rotates and the stroke reverses, such that the suction surface of the airfoil switches sides (Fig. 5B).

**Figure 4.**
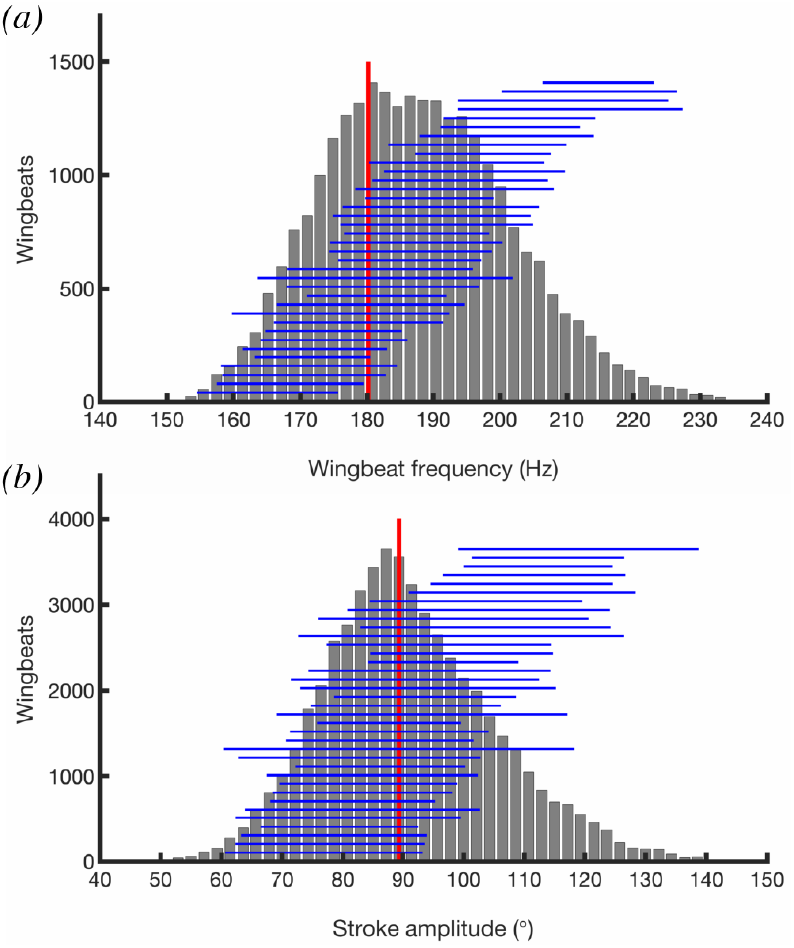
Histograms of summary parameters for the wing kinematics. *(a)* Wingbeat frequency. *(b)* Stroke amplitude. The vertical red line shows the kinematics of the standard hovering wingbeat. Horizontal blue lines span the 5^th^ to 95^th^ percentiles for each individual, ranked according to their mean. Note that most of the variation in these parameters is seen between, rather than within, individuals.

**Figure 5.**
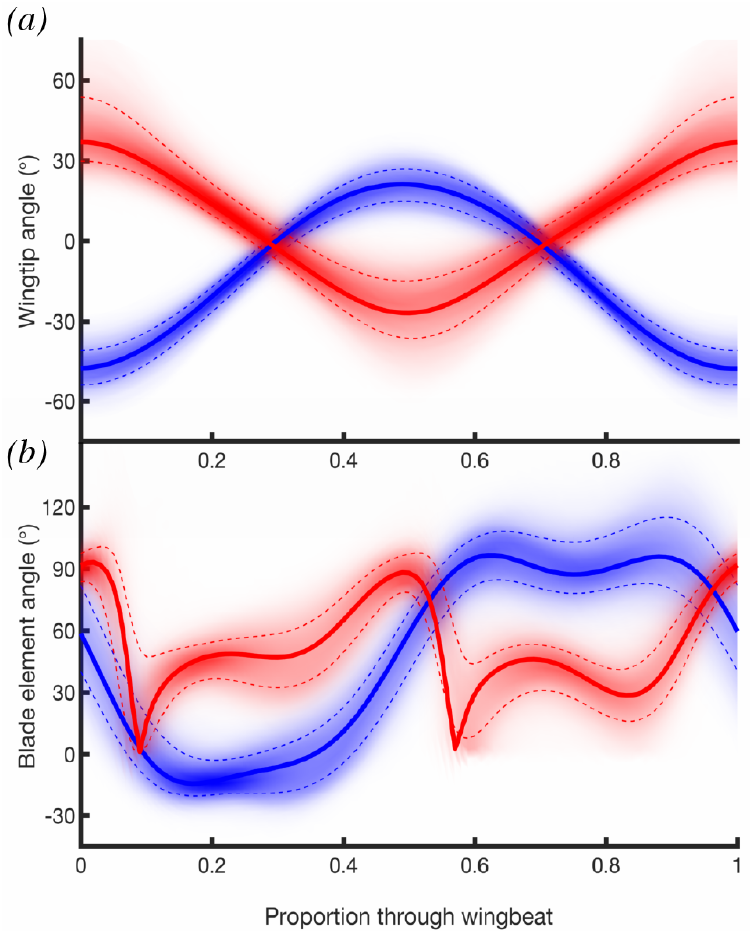
Frequency density plots of time-varying wing kinematics over all wingbeats. *(a)* Wing tip kinematics, showing stroke angle *ϕ* (red), and deviation angle *θ* (blue). *(b)* Blade element kinematics, showing wing pitch angle *ω* (blue), and aerodynamic angle of attack *α* (red), both measured at midspan. The shading density corresponds to the frequency density, conditional upon wingbeat phase. Dashed lines indicate ±1SD from the mean; solid lines plot the kinematics of the standard hovering wingbeat.

### (c) Aerodynamic force coefficients

With the aerodynamic speed *U* and angle of attack *α* defined at the three-quarter chord point, our best estimates for the model parameters were 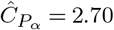 and 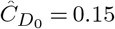, such that the lift and drag coefficients (Eqs. 2.10 and 2.12) may be modelled empirically as:

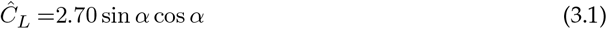

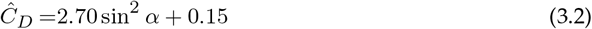

The variance in the estimated lift and drag curves over 10^5^ random 10% subsamples of the 854 flight sequences was negligible for the lift coefficient but more substantial for the drag coefficient (Fig 6B). This reflects the fact that the error in the estimation of the lift coefficient depends only on the error in the estimation of the aerodynamic force coefficient derivative 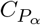, whereas error in the estimation of the drag coefficient depends also on the error in the estimation of the drag coefficient offset 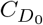. We therefore investigated the effect of dropping 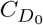 from the regression model in Eq. 2.20 (i.e. modelling the drag coefficient using Eq. 2.11 instead of Eq. 2.12). This produced a 1% increase in the estimated aerodynamic force coefficient derivative 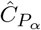, from 2.70 to 2.73, in compensation for the slight decrease in the predicted drag that would otherwise result from dropping 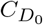. These changes were associated with only a 0.6% increase in the error sum of squares (Table 1), so the inclusion of a drag coefficient offset—though justified theoretically—adds little predictive power to the model. On the other hand, it is clear from Fig 6B that the estimated value of the drag coefficient at zero angle of attack is consistently positive across many different subsamples of the data, so the inclusion of 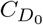 in the model is also justified empirically.

**Figure 6.**
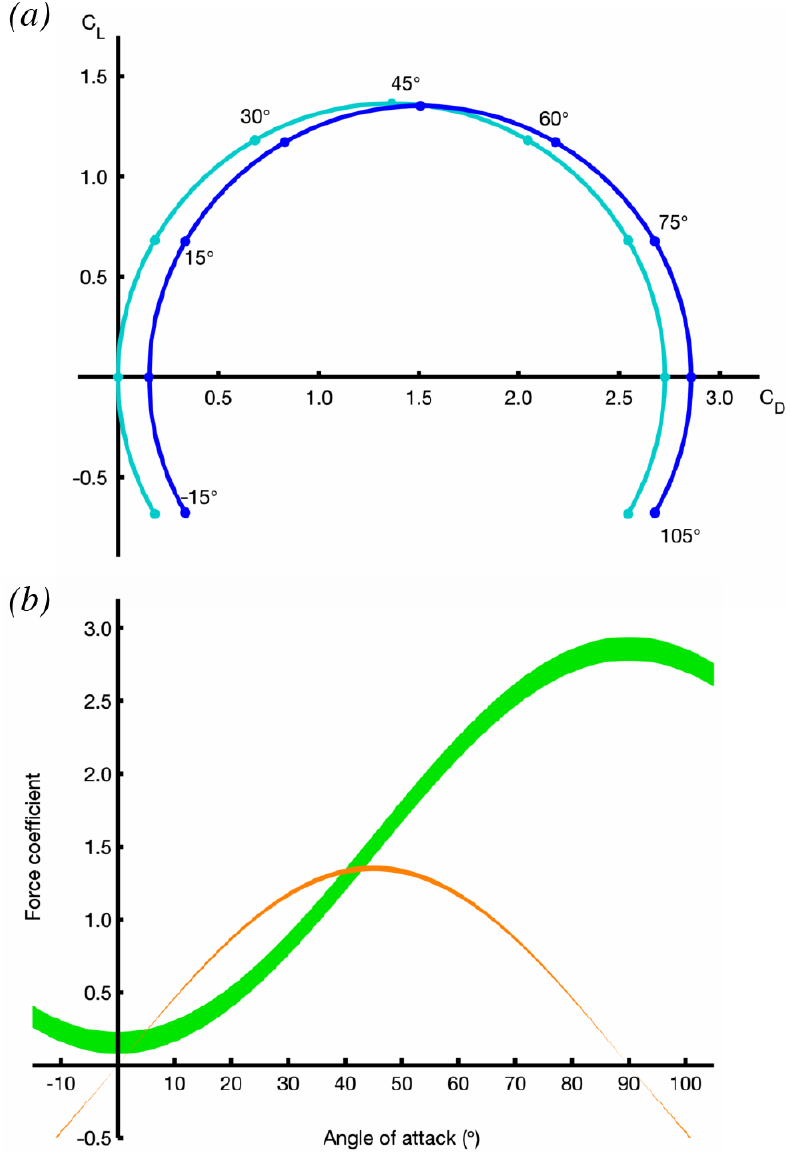
Empirically-fitted lift and drag polars. *(a)* Fitted lift coefficient *Ĉ*_*L*_ versus fitted drag coefficient *Ĉ*_*D*_ across the full range of angles of attack *α*, for the full model with two empirically-fitted parameters (blue), and for the reduced model with no drag offset term (cyan). *(b)* Fitted lift coefficient *Ĉ*_*L*_ (orange) and fitted drag coefficient *Ĉ*_*D*_ (green) plotted against angle of attack *α*. The width of the lines indicates their mean ±1SD, assessed over 10^5^ random subsamples of the data, each comprising 10% of the recorded flight sequences.

These results assume that the aerodynamic speed *U* and angle of attack *α* are defined at the three-quarter chord point, which is a classical result of lifting-line theory and unsteady airfoil theory in the limit of small amplitude motion (see Section 2(e)). Adjusting the reference point at which the kinematics were defined allowed a small reduction in the error sum of squares, but with the paradoxical result that the estimated drag coefficient offset 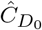 became negative if the reference point was moved further than 85% towards the trailing edge (Fig. 7). Subject to the physical constraint that 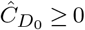, the error sum of squares was minimised when *U* and *α* were defined at 87% of the chordwise distance back from the leading edge, with 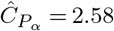 and 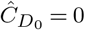. The chordwise reference point minimising the error sum of squares is therefore associated with a binding inequality constraint because 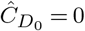 at this optimum. Since this optimisation produces at best a 1% improvement in the error sum of squares and comes at the cost of estimating a third parameter from the data using an exhaustive search procedure (Table 1), we prefer to retain the parameter estimates of 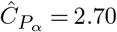 and 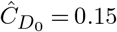 with the kinematics defined at the three-quarter chord point, in accordance with our prior expectation from classical aerodynamic theory (see Section 2(e)).

**Figure 7.**
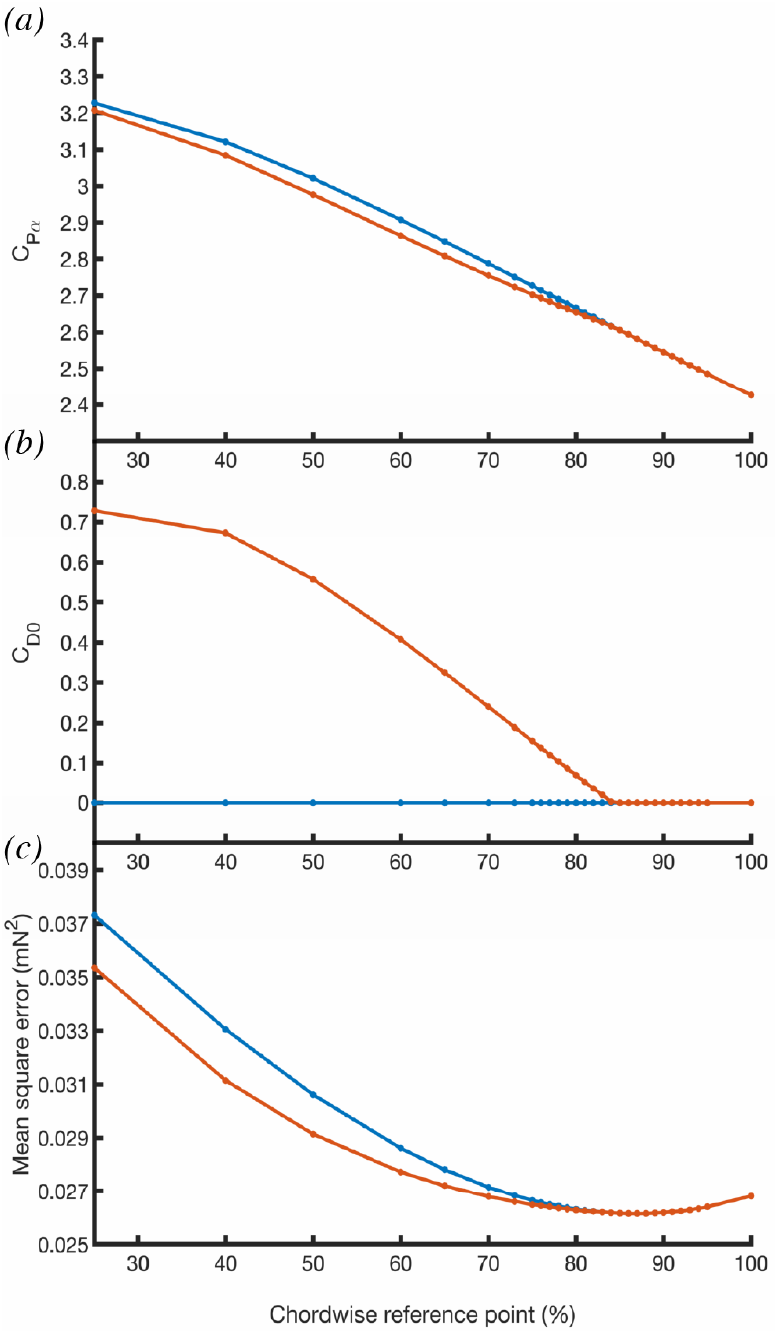
Variation in the parameter estimates of the blade element model as a function of the chordwise position of the reference point at which the aerodynamic speed *U* and angle of attack *α* are defined: (A) estimated pressure force coefficient derivative, 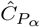 ; (B) estimated drag coefficient offset, 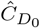 ; (C) associated mean squared error (MSE); see Table 1 legend for details of calculation. These parameter estimates are made subject to the inequality constraint that 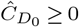, which bites when the kinematics are defined at *≥* 85% chord. See text for further details.

### (d) Goodness of fit of the stroke-averaged aerodynamic forces

Because the blade element model is fitted as a regression forced through the origin (Eq. 2.20), its *R*^2^ statistic is not well defined. To assess its goodness of fit with respect to the measured forces, we therefore regressed the fitted aerodynamic forces on the measured aerodynamic forces, without forcing the regression line through the origin (Fig. 8). We did this separately for each body axis, and found that a large proportion of the variation in the stroke-averaged forces measured in the body’s plane of symmetry was explained in both the *x*_*b*_- and the *z*_*b*_-axes (*R*^2^ = 82.5% and *R*^2^ = 79.0%, respectively). In contrast, a much smaller proportion of the measured variation in the stroke-averaged forces was explained in the transverse *y*_*b*_-axis (*R*^2^ = 18.0%). Moreover, although the regression intercept was appropriately close to zero in all three body axes (*x*_*b*_: 0.041 mN; *y*_*b*_: 0.000 mN; *z*_*b*_: 0.021 mN), the regression slope was only suitably close to one in the body’s plane of symmetry (*x*_*b*_: 0.92; *z*_*b*_: 1.05), being greatly attenuated by noise in the transverse axis (*y*_*b*_: 0.62). This poor fit in the *y*_*b*_-axis presumably reflects the fact that the total range of the lateral stroke-averaged aerodynamic forces was small (Fig. 3e), which leads to a lower signal to noise ratio in the *y*_*b*_-axis (Fig. 8B) than in the *x*_*b*_- or *z*_*b*_-axes (Fig. 8A,C). The forces measured in the *x*_*b*_- and *z*_*b*_-axes are generally well modelled, although a detailed inspection of the regression plots shows that whereas the aerodynamic forces measured in the *x*_*b*_-axis are fitted closely over their entire range (Fig. 8A), the blade element model systematically under-predicts the magnitude of the largest aerodynamic forces produced in the *z*_*b*_-axis (bottom left region of Fig. 8C). Even so, the blade element model does a good job of fitting the overall time-history of the measured stroke-averaged forces on the timescale of an entire flight sequence (Fig. 9).

**Figure 8.**
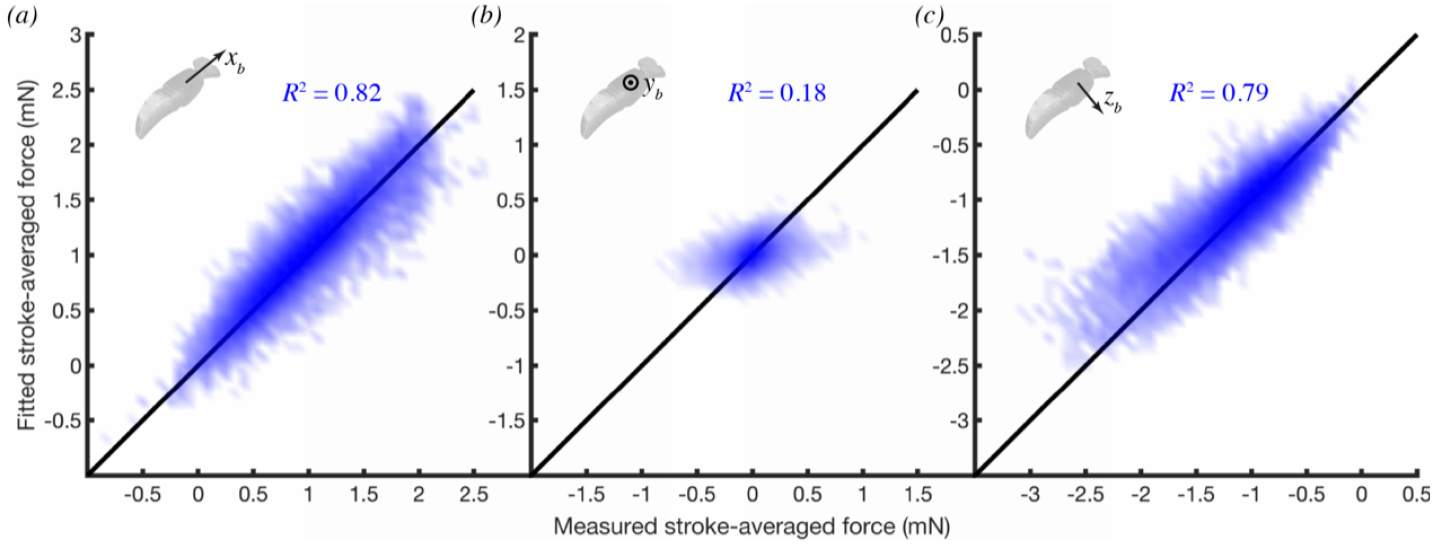
Two-dimensional histograms of fitted versus measured stroke-averaged forces for all wingbeats, resolved in the three body axes *(a-c)*. Shading density corresponds to frequency density of data; black line indicates the ideal line of slope one, passing through the origin. Note that at peak aerodynamic force production, the regression model systematically under-predicts the magnitude of the force actually produced in the *z*_*b*_ axis.

**Figure 9.**
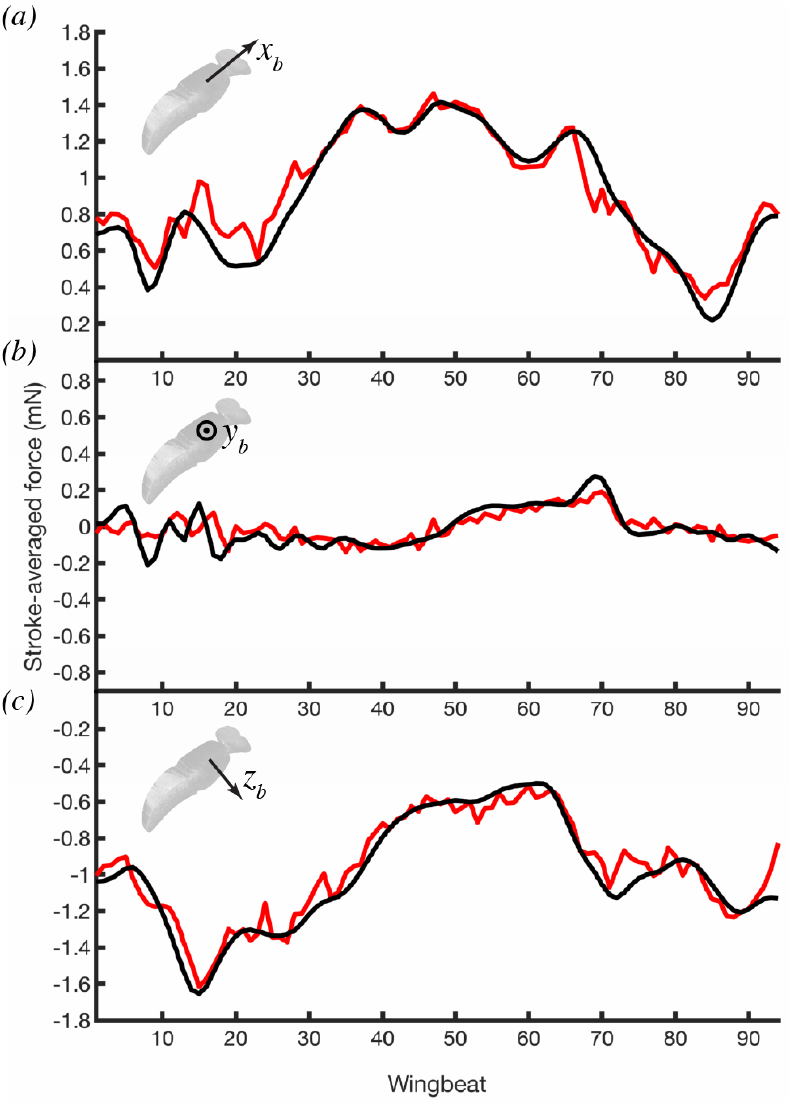
Time history of the measured and fitted stroke-averaged aerodynamic forces for one flight sequence, resolved in the three body axes *(a-c)*. The forces fitted by the model (red) show a close match to the measured forces (black) in all three body axes. See Supplementary Video S1 for the wing and body kinematics corresponding to this flight sequence.

### (e) Predicted aerodynamic forces through the wingbeat

Because the parameters of the blade-element model were fitted only to the stroke-averaged forces, there is no necessary statistical reason to assume that the resulting model will perform well in predicting the time-varying aerodynamic forces through the wingbeat, but the physical basis of the underlying aerodynamic model is such that it could be expected to. We used the blade element model to predict how the aerodynamic forces are expected to vary through the standard hovering wingbeat that we defined in Section 2(b) (Fig. 10, 11; Supplementary Videos S2, S3), to allow us to assess the relative contributions of lift, drag, and added mass at different stages of the wingbeat. As a further check on the robustness of our predictions to errors in parameter estimation, we modelled the time-varying aerodynamic forces through this standard hovering wingbeat, across the full range of variation in the aerodynamic force coefficients estimated for the subsamples in Section3(c). Despite the variation in the aerodynamic force coefficient parameters fitted in the subsampling analysis (Fig. 6B), the resulting variation in the predicted time-varying aerodynamic forces was slight in comparison to their variation through the wingbeat (Fig. 10). The results in Fig. 10 therefore provide a sound basis for comparing the predictions of our blade element model with future CFD simulations of the standard hovering wingbeat.

**Figure 10.**
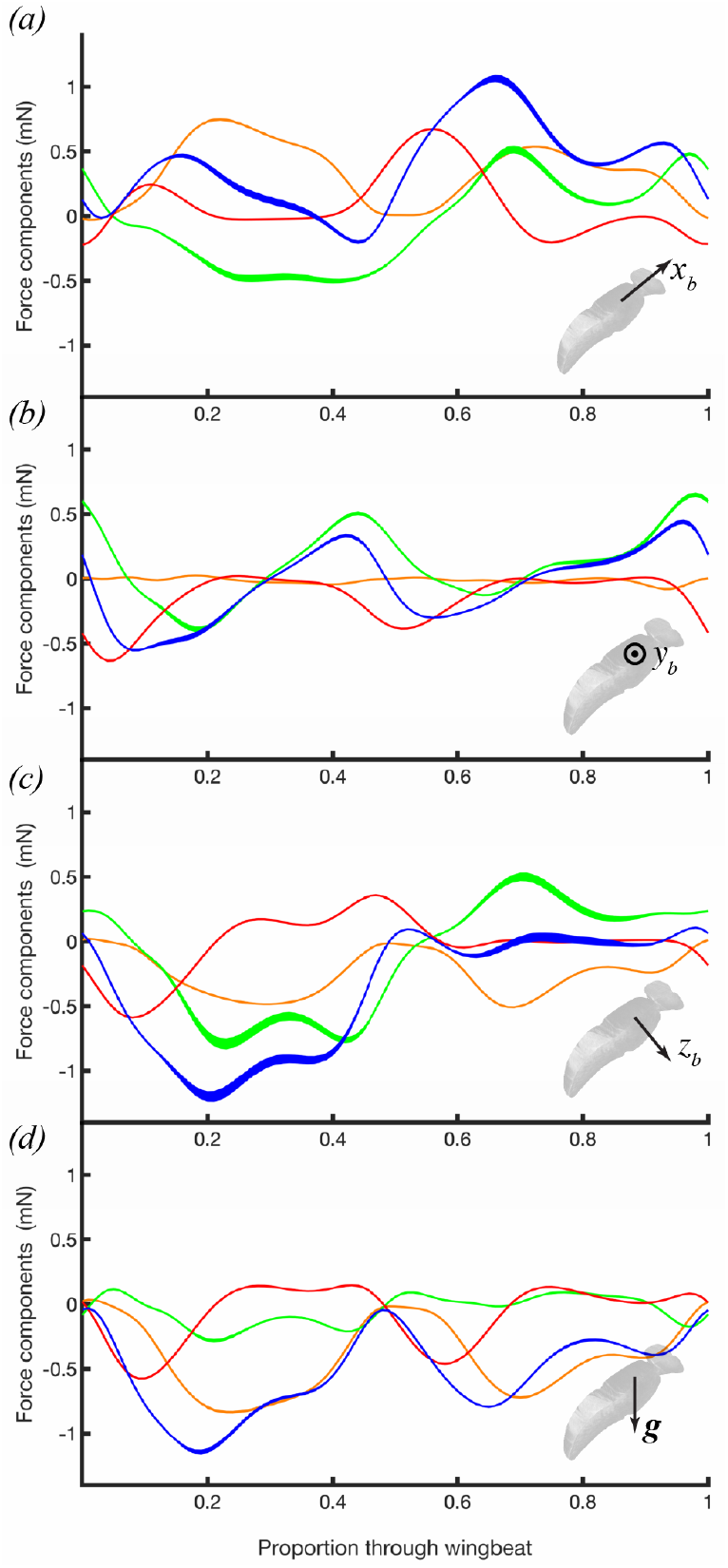
Predicted time-varying force components for the standard hovering wingbeat. The plots show the forces predicted for the right wing only, decomposed as lift (orange), drag (green), and added mass (red), together with their sum (blue). (*a-c*) Forces resolved in the body axes. (*d*) Vertical aerodynamic force component. The width of the lines indicates their mean ±1SD, assessed over 10^5^ random subsamples of the data, each comprising 10% of the recorded flight sequences.

**Figure 11.**
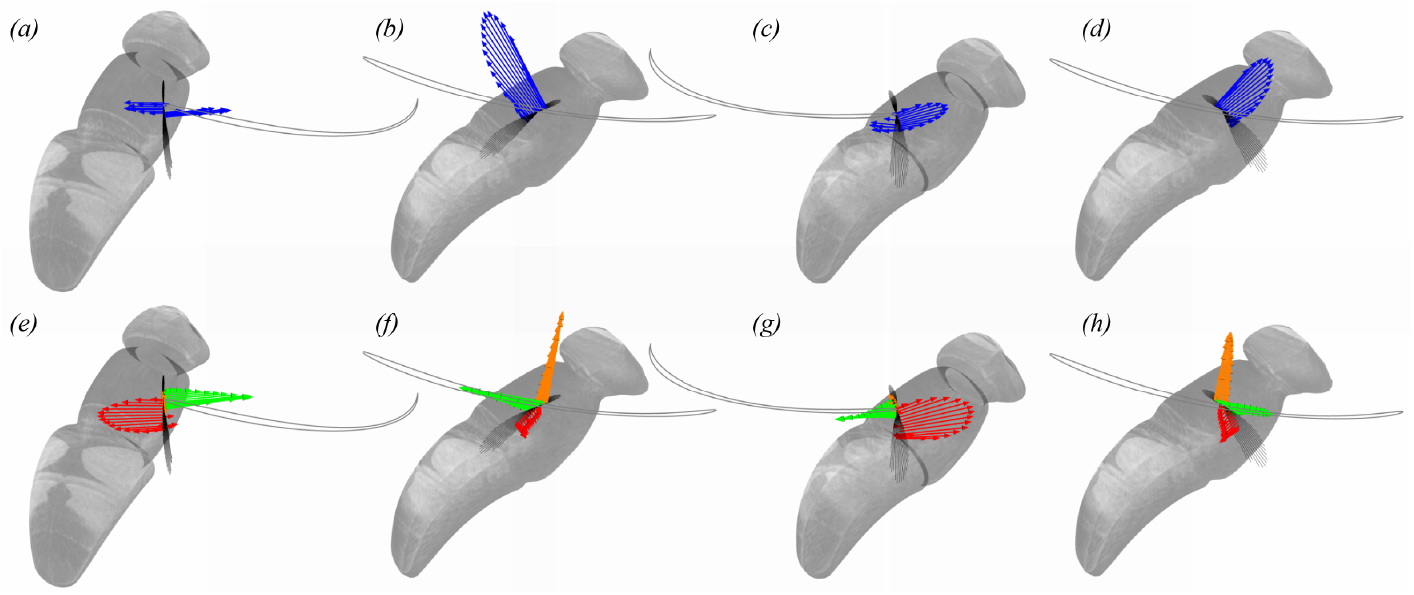
Spanwise distribution of predicted aerodynamic force at four phases of the standard hovering wingbeat. (*a-d*) Resultant aerodynamic force (blue arrows). (*e-h*) Quasi-steady lift (orange arrows) and drag (green arrows) force, together with unsteady added mass force (red). The four phases of the wingbeat correspond to: (*a,e*) beginning of the downstroke; (*b,f*) middle of the downstroke; (*c,g*) beginning of the upstroke; (*d,h*) middle of the upstroke. The hoverfly body is shown in the background, and the grey line shows the wing tip path. The viewpoint rotates with the stroke, so as to look down the spanwise axis of the wing at all times. Individual blade elements are plotted as black lines, to show the spanwise distribution of wing twist through the wingbeat. See Supplementary Videos S2 and S3 for animated versions of these forces at all time steps.

Because the stroke plane is close to horizontal during the standard hovering wingbeat, weight support is attributable primarily to aerodynamic lift, which is predicted to account for 77.1% of the stroke-averaged vertical force (Fig. 10D). The added mass and drag forces each make a small net positive contribution to weight support, providing 14.6% and 8.3%, respectively, of the predicted stroke-averaged vertical force. The predicted lift force peaks at the middle of each half-stroke, when the wing’s translational velocity is highest (Fig. 11F,H; Supplementary Video S3), so this is also the phase of the stroke during which the majority of the vertical force is expected to be produced (Fig. 10D; Fig. 11B,D). In contrast, the added mass force peaks at the beginning of each half-stroke, when the wing’s acceleration normal to its chord is highest (Fig. 11E,G; Supplementary Video S3). The drag force has a more complicated time-history again, peaking at several points through the stroke (Fig. 11E-H; Supplementary Video S3). Aerodynamic force production in the transverse *y*_*b*_-axis is qualitatively similar on both the upstroke and downstroke (Fig. 10B), and the same is true of the vertical component of the aerodynamic force (Fig. 10D), but in each case the amplitude of the forces is somewhat diminished on the upstroke relative to the downstroke. In contrast, aerodynamic force production along the *x*_*b*_- and *z*_*b*_-axes of the body displays a marked asymmetry between the upstroke and the downstroke (Fig. 10A,C). The dynamics of hovering force production are therefore considerably more complex than might first appear from the time-history of the vertical aerodynamic force component alone (Fig. 10D).

## 4. Discussion

Perhaps the greatest challenge in modelling insect flight is to predict the aerodynamic forces that the flapping wings impart as they undergo a variety of complex aeroelastic motions in a variety of different flight conditions. Although current CFD techniques allow these forces to be predicted with great accuracy for a given set of wing or body kinematics, it remains extremely time consuming to compute the flows associated with variable wing or body kinematics or with different wing morphologies [8,16,31,42]. Moreover, the aerodynamic assumptions that classical analytical models must make to fix a solution for the aerodynamic force coefficients are so restrictive as to prevent their realistic application to insect flight [11]. Here we have aimed to find a middle ground, by estimating the aerodynamic force coefficients empirically for a simple analytical blade element model that captures the scaling of the forces expected from first principles, and which models the forces with sufficient accuracy to be used in a range of other applications. By estimating just two numerical parameters—the derivative of the pressure force coefficient 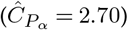 and the drag coefficient offset 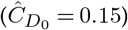 —for the largest dataset of insect wing and body kinematics obtained to date [39], we have captured 80% of the total variation in the measured stroke-averaged aerodynamic forces in the sagittal plane over *N* = 26, 541 wingbeat pairs recorded in freely hovering and manoeuvring *Eristalis* hoverflies. This is possible because whereas the aerodynamic form of the model is quite simple and uses empirical estimates of the force coefficients that are averaged over many different flight conditions, the modelling of the kinematics is sufficient to capture almost the full complexity of the wingbeat. In summary, the key contribution of this paper is to provide a kinematically accurate blade-element model of the stroke-averaged forces of insect flight, fitted and validated with respect to an extensive free-flight dataset including a wide range of flight manoeuvres.

### (a) Key features of the modelling

A key feature of our kinematic model is that it takes full account of the torsional deformation of the wing, without either limiting the motion to a fixed stroke plane, or treating the wing as a flat plate. This matters, because wing flexion is a defining characteristic of insect flight, which can reduce its aerodynamic power requirements and enhance the useful force produced [23,42]. For example, when we tried treating the wing as a flat plate operating at a pitch angle matched to that of the twisted wing at mid-span, we found that the error sum of squares increased by 30%, which was associated with a large decrease in the estimated value of 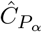 and an unrealistically large increase in the estimated value of 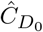 (Table 1; Fig. S3c). Modelling the kinematics accurately is therefore at least as important as modelling the flow accurately, so it does not make sense, for example, to go the effort of solving the full Navier-Stokes equations numerically for an oversimplified model of an insect’s wing kinematics.

Another key feature of the kinematic modelling is that it implicitly accounts for the body’s own velocity and acceleration when computing the velocity, angle of attack, and acceleration of each blade element. The body’s motion plays a key role in flight stability [e.g. 15,32], so it is essential that any aerodynamic model which aims to investigate free-flight behaviour takes account of these effects. Moreover, whereas the velocity of the body is usually considerably smaller than the velocity of the wing tip, the two can become comparable in magnitude during fast manoeuvres and fast forward flight. In fact, there is clear evidence of the importance of the body’s motion in our modelling, because if we ignored the body’s velocity and acceleration when fitting the blade element model, then the error sum of squares increased by 34%, and the estimate of the drag coefficient offset became unreasonably high (Table 1; Fig. S3d).

Concerning the aerodynamic modelling, it is noteworthy that moving the reference point at which the kinematics are defined backwards from its default three-quarter chord position results in at best a 1% reduction in the error sum of squares which is associated with the drag coefficient offset being driven unrealistically towards zero. Conversely, moving the reference point forwards from the three-quarter chord point causes a rapid worsening of the model fit, which implies that the default approach is successfully capturing the rotational lift as expected (see Section 2(e)). This provides a useful empirical validation of the approach of treating rotational lift together with translational lift [35] by defining the aerodynamic speed *U* and angle of attack *α* at the three-quarter chord point [11], consistent with the results of previous work using mechanical flappers [31].

### (b) Comparison to other models

Blade element models have been used previously to predict the aerodynamic forces of insect flight, but never in combination with such detailed kinematic data from free-flying insects. Our model differs from others used previously in the following respects: (i) it includes the full three-dimensional motion of the wing, including torsional deformation; (ii) it incorporates all six rotational and translational degrees of freedom of the body; (iii) it models the rotational aerodynamic force by calculating the angle of attack at the three-quarter chord point rather than treating this as a separate contribution; (iv) it captures systematic variation in the direction of the quasi-steady aerodynamic force with respect to the wing’s surface and the flow; and (v) it has a simple and transparent aerodynamic form developed from physical principles. The first two of these distinguishing features relate partly to the availability of data, but all are fundamental.

Concerning our treatment of the rotational forces, most recent blade element models include a separate rotational lift component, sometimes called a rotational circulation force [e.g. 31,34]. Indeed, the blade element model of Nakata *et al*. [26], with coefficients fitted using computational fluid dynamics also includes a separate rotational drag force. These rotational forces are expressly intended to capture the aerodynamic effects of pitching or twisting about the spanwise axis of a flapping wing, but by defining the velocity of each blade element at its three-quarter chord point [11], all or most of these rotational effects can be incorporated directly into the calculations of translational lift and drag, thereby simplifying the model conceptually and reducing the number of free parameters that must be estimated [35].

The direction of the quasi-steady pressure force is determined in our model by the balance of the orthogonal lift and drag forces, where the lift on each blade element is assumed to act perpendicular to the relative air flow, and where the drag is assumed to act in the direction of the relative air flow. Other blade element models have defined the circulatory force as acting normal to the blade element, but have assumed that its magnitude is equal to that of the resultant lift and drag [16,30,31,34], which means that a circulatory force is assumed to act perpendicular to the chord even at zero angle of attack if there is any friction drag or pressure drag under these conditions. In contrast, the treatment of the drag offset term in our model means that the direction of the quasi-steady aerodynamic force varies with respect to the chord (Fig. 12A). Specifically, whereas the quasi-steady force is almost perpendicular to the chord at angles of attack greater than about 45°, it becomes tilted back substantially at lower angles of attack, ultimately becoming tangent to the chord at zero angle of attack.

**Figure 12.**
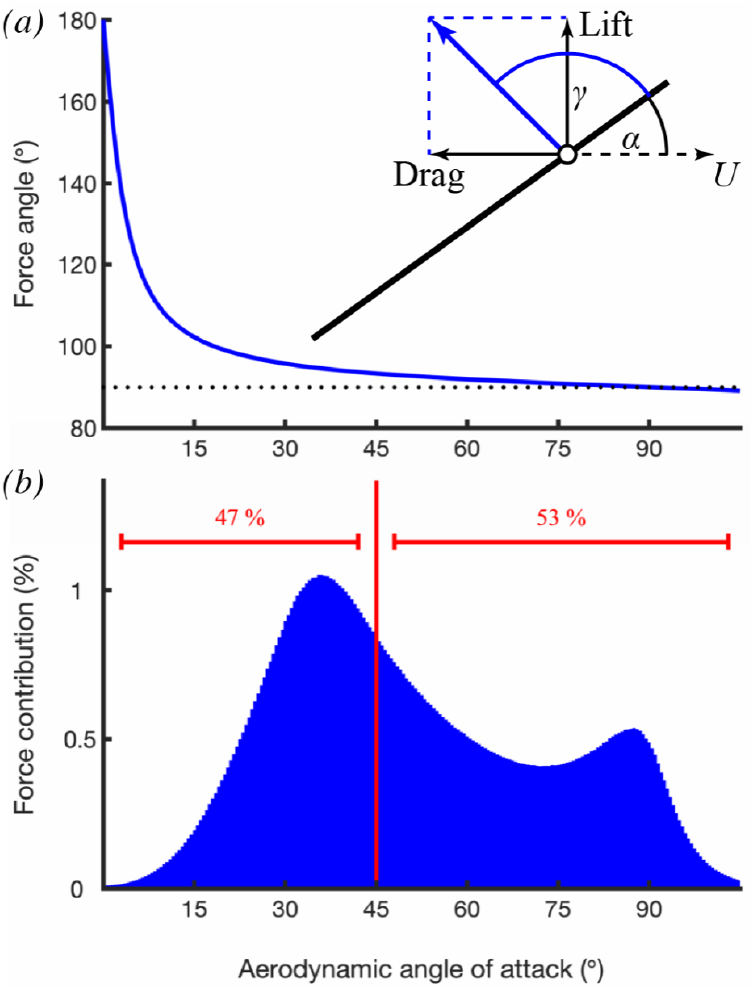
Predicted direction and relative contribution of the quasi-steady aerodynamic force across different angles of attack. (*a*) The direction of the resultant quasi-steady force vector is represented by its angle (*γ*) with respect to the blade element chord, across the full range of measured aerodynamic angle of attack (*α*). The predicted angle *γ* is calculated as the angle between the resultant lift and drag force and the blade element (inset diagram), using the regression estimates for 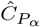 and 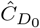. This angle *γ* only approaches 90° (dotted line) at angles of attack above approximately 45°. (*b*) Distribution of predicted contribution to the stroke-averaged quasi-steady aerodynamic force at different aerodynamic angles of attack for all recorded wingbeats combined. The horizontal red bars display the total percentage of the force produced at angles of attack above and below 45° (i.e. the area under the graph to either side of the vertical red line).

This behaviour is appropriate given the inevitable presence of friction drag, and the likely importance of pressure drag even at low angles of attack. Although insect wings are commonly treated as approximating an idealised thin aerofoil, they do in fact have a finite thickness, particularly at the leading edge, which typically functions as a reinforced spar. Wing corrugation due to venation also makes wings inherently three-dimensional structures, which can further increase profile drag [36,38]. Aerodynamic measurements of real insect wings [9,10], or mechanical and computational models thereof [6,28], have all indicated the presence of significant drag at zero angle of attack, and hence of the tilting back of the resultant aerodynamic force at low angles of attack. Whilst it is true that insect wings operate at characteristically high angles of attack, in the hoverfly data that we have presented here, 47% of all of the aerodynamic force produced by the wing is predicted to occur at angles of attack *α* < 45° (Fig. 12B), with a mode at *α* = 36°, for which the force vector will be tilted back significantly from 90° (Fig. 12A).

Perhaps the most important feature of our model is the simplicity of its form, which despite being highly nonlinear in the kinematics, is nevertheless linear in its two free parameters 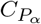 and 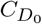. This means that we are able to fit these parameters analytically using linear least squares, guaranteeing that they are optimised globally and at speed—even on the very large dataset that we employ. Moreover, because the form of our model was developed from first principles rather than with reference to the data, it provides useful physical insight into the *ad hoc* form of some influential models that have been fitted previously. In particular, Dickinson *et al*. [6] modelled the lift and drag coefficients for their robotic flapper as *C*_*L*_ = 0.225 + 1.58 sin (2.13*α* − 0.14) and *C*_*D*_ = 1.92 − 1.55 cos (2.04*α* − 0.17), where *α* is in radians and where all of the numerical constants are estimated from the data. These complicated formulae use a total of eight free numerical parameters to model *C*_*L*_ and *C*_*D*_, but make better sense physically when it is observed that they approximate trigonometric double angle formulae. Noting that sin (2*α*) = 2 sin *α* cos *α* and cos (2*α*) = 1 − 2 sin^2^ *α*, Eqs. 3.1 and 3.2 describing our own fitted model can be restated as:

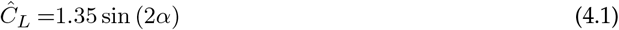

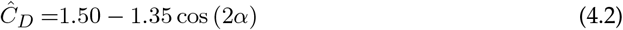

which are not dissimilar to the equations fitted by Dickinson *et al*. [6] that themselves approximate trigonometric double angle formulae. This in turn reflects the fact that whereas the resultant force varies sinusoidally with the angle of attack, its direction is approximately normal to the wing’s surface, such that its lift and drag components resolved perpendicular and tangent to the relative airflow vary as cosine and sine functions of the angle of attack, respectively.

### (c) Extension to other datasets

The same blade element model can be straightforwardly applied to other insect species, because other than the morphological parameters of wing length and wing shape (Fig. 2), there are no modelling assumptions that are specific either to hoverflies or to the dataset that we used. Indeed, one of the strengths of our approach is that the same blade element model can be straightforwardly fitted to any similar data set, using linear least squares to optimise the force coefficient parameters: Matlab code implementing the blade element model is provided as Supporting Data S1 to this end. It is clear from our modelling that knowledge of the wing twist distribution is essential to fitting the forces accurately (Table 1), which reinforces the need for future kinematic studies to measure wing deformation as we have done here. Of course, in design problems where the wing twist distribution is a parameter that may need to be optimised rather than measured, the inclusion of wing twist in our analytical blade element model makes it suitable for use in fast global optimization of the wing deformation parameters prior to local refinement using CFD.

Our analysis shows that the blade element model is robust to drastic reductions in sample size, with data subsampling producing comparatively small changes in the predicted forces when fitting the force coefficients to random subsamples comprising only 10% of the recorded data. This means that the same modelling approach can be applied to datasets much smaller than the *N* = 26, 541 wingbeats that we analyse here. If necessary, the simplicity of the model can be increased further by excluding the small drag offset term. This then means that only the derivative of the pressure force coefficient need be estimated, which reduces the variance of the parameter estimate (Fig. S3b). Given the comparatively weak signature of the drag offset term at high angles of attack, this may be preferable approach when working with smaller or noisier datasets than the one available here. However, whilst the inclusion of the drag offset term provides only a 0.6% reduction in the error sum of squares, we retain it in our model because its importance is supported by both theory and experiment [6,28]. Furthermore, although its inclusion has a minimal effect on the accuracy of our modelling of this dataset for *Eristalis*, the drag offset term may be more important for other species with different morphologies and especially for those operating at lower Reynolds numbers.

As with any form of regression modelling, an important caveat is that the dataset must contain sufficient variation in the independent variables to enable a good fit (compare, Fig. 8a,c with Fig. 8b). In principle, our regression estimates of the lift and drag coefficients could be replaced by estimates from model wings [6] or CFD [26]. However, an obvious risk of this approach is that the aerodynamic properties of a model wing, or even those of a detached wing suffering rapid desiccation [22], may differ markedly from the aerodynamic properties of a real wing *in vivo*. These problems are avoided completely by our approach of fitting the parameters of the blade element model empirically to free-flight data from live insects.

### (d) Limitations

Although our blade element model fits the aerodynamic forces well in the sagittal plane over most of their range, it systematically under-predicts the magnitude of the largest aerodynamic forces produced in the *z*_*b*_-axis (Fig. 8C). This discrepancy presumably indicates a nonlinearity in aerodynamic force production that the quasi-steady blade element model fails to capture. More specifically, as most of the force in the *z*_*b*_-axis is produced on the downstroke (Fig. 10C), we hypothesise that this non-linearity reflects some unsteady aerodynamic mechanism that becomes increasingly important as force production is increased on the downstroke. One obvious possibility is that this non-linearity is due to the presence of a leading-edge vortex (LEV) on the functional upper surface of the wing. This is one of the characteristic aerodynamic mechanisms of insect flight, allowing the lift curve to be extended to high angles of attack by delaying stall [12]. Delayed stall is already implicit in our model because of its explicit assumption that the wing does not suffer a sudden loss of lift at high angles of attack (Fig. 6). It remains unclear, however, whether the presence of an LEV enhances the aerodynamic force coefficients by amplifying some portion of the lift curve rather than merely by extending it [24]. Hence, although the average aerodynamic effect of the LEV should be captured by the empirical force coefficients that we have estimated from our data, it is plausible that the model might still underestimate the lift enhancement provided by the LEV at very high angles of attack. It is also worth noting that our model does not account explicitly for the three-dimensional effects of spanwise flow impacting the strength and stability of the LEV [27], nor for the possible effects of interactions between the wing and the wake shed on the preceding half-stroke [31].

A second possibility is the clap-and-fling mechanism [11], which can occur if the wings approach one another closely at the top of the upstroke and are then flung apart on the downstroke (Fig. 13A). This mechanism may also be modified by the effects of spanwise bending and chordwise camber [18], which we do not model directly here. Interestingly, although there is a positive association between the stroke amplitude and the magnitude of the measured aerodynamic force along both the *x*_*b*_- and *z*_*b*_-axes (Fig. 13B), as expected under a quasi-steady model of the forces, this association is much stronger in *x*_*b*_ (*R*^2^ = 0.53) than in *z*_*b*_ (*R*^2^ = 0.27). This suggests that increases in the magnitude of the aerodynamic forces in *z*_*b*_ are not principally driven by increases in stroke amplitude. On the other hand, there is a negative association between the wing tip separation at the start of the downstroke, and the magnitude of the measured aerodynamic forces (Fig. 13C), which is stronger in the *z*_*b*_-axis (*R*^2^ = 0.62) than in the *x*_*b*_-axis (*R*^2^ = 0.25). This negative association would be expected under an unsteady clap-and-fling mechanism, and the strength of the association in *z*_*b*_ suggests that increases in the magnitude of the aerodynamic forces in this axis may indeed by driven by an unsteady clap-and-fling mechanism. This is consistent with the interpretation that wing-wing interactions affect force production in the *z*_*b*_-axis more than in the *x*_*b*_-axis. Since the blade element model will not capture this nonlinearity directly, it follows that the linear parameter estimate for 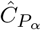 can be expected to overestimate the true value of the quasi-steady force coefficient derivative 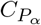.

**Figure 13.**
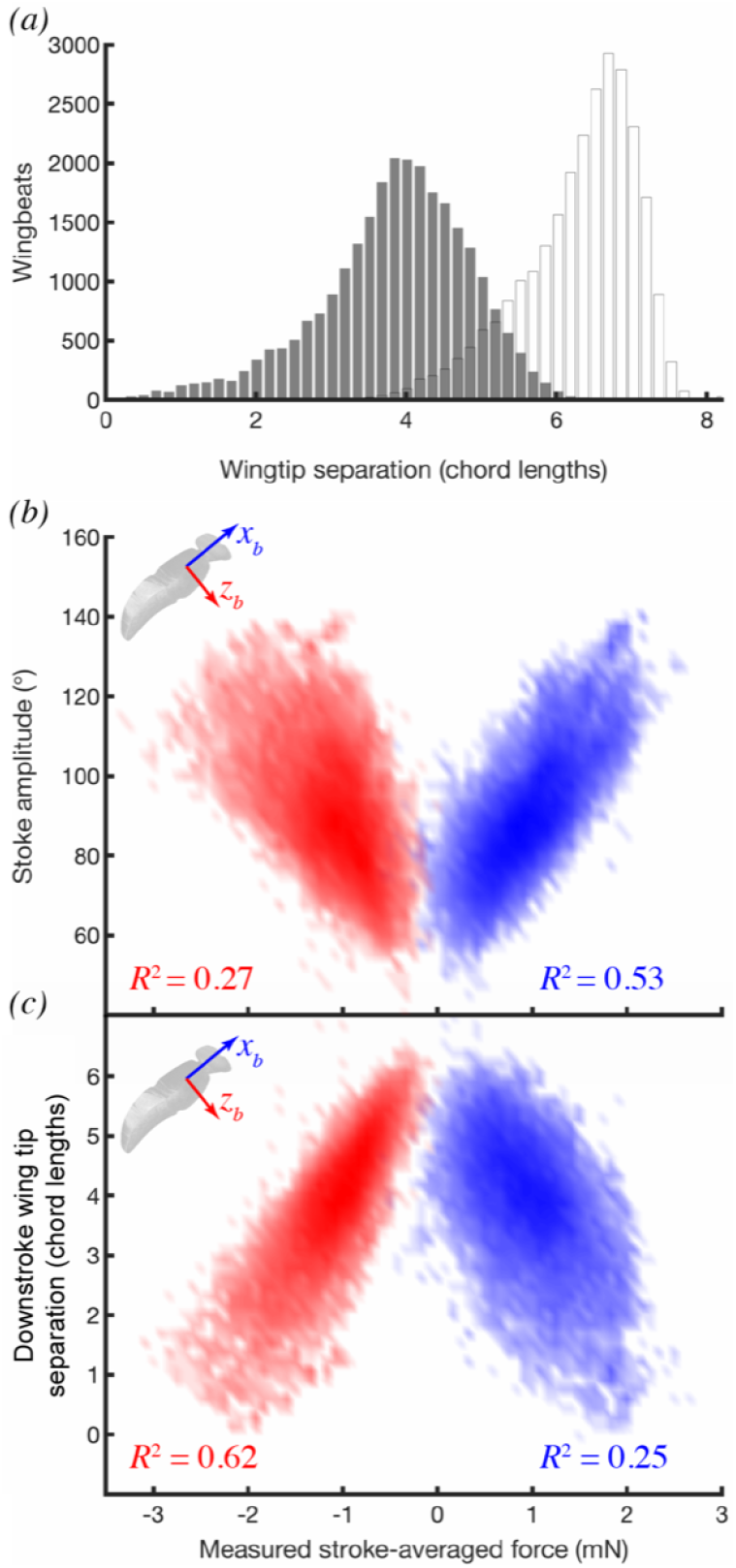
Histograms of wing tip separation and stroke amplitude, and their association with the measured stroke-averaged aerodynamic force. *(a)* Histograms of wing tip separation at the start of the downstroke (shaded bars) and at the start of the upstroke (unshaded bars). *(b,c)*, Frequency density plots showing stroke amplitude and wing tip separation at the start of the downstoke, versus the measured forces in the *x*_*b*_-(blue) and *z*_*b*_- (red) axes of the body. Shading density corresponds to frequency density of data. Wing tip separation was calculated as the distance between the wing tips at the start of the half-stroke, normalised by the mean wing chord; *R*^2^ statistics for the linear associations between the wingbeat parameters and the measured forces are shown for each axis.

Finally, although the effects of body motion are captured in our modelling of the wing kinematics and aerodynamics, it is important to note the body itself will produce drag—and perhaps some lift—in forward flight [10]. Modelling the aerodynamic forces produced by a bluff body is not straightforward, owing to the likelihood of sudden flow separation above some critical angle of attack, but as most of the flight sequences that we modelled were close to hover, it is reasonable to assume that the aerodynamic forces on the body would have been overwhelmed by the aerodynamic forces acting on the wings at most stages of the wingbeat.

## 5. Conclusions

We have shown here that an analytical blade element model with just two empirically-fitted coefficients provides a close fit to the measured stroke-averaged aerodynamic forces of free-flying insects in manoeuvring flight. The alternative approach of using computational fluid dynamics modelling is capable of capturing fine aerodynamic detail, and has even led to the discovery recently of novel unsteady mechanisms [e.g. 4], but is computationally expensive, taking many orders of magnitude longer to deliver results than the analytical blade element model presented here. Both approaches therefore have a complementary role to play. Analytical blade element modelling functions well for investigating large datasets, studying the effect of changing wing kinematics parametrically, and making quick comparisons across species. Conversely, a numerical approach is preferable where high-fidelity predictions, detailed time histories, or insight into unsteady aerodynamic mechanisms is required. The strengths of the analytical blade element model that we have presented here are: (i) the simplicity of its underlying aerodynamic equations; the complexity of the deforming wing kinematics and body motions that it models; and (iii) the fact that its aerodynamic force coefficients are fitted empirically to free-flight data from real insects, thereby capturing the full scope of the insect’s flight dynamics. Besides demonstrating the importance of prioritising accurate modelling of the deforming wing kinematics ahead of detailed modelling of the fluid dynamics, we expect that our model will serve as a useful, validated tool for future research on insect flight dynamics and control.

## Data Accessibility

The supplementary data and code supporting this article are available through figshare https://doi.org/10.6084/m9.figshare.11725929.

## Authors’ Contributions

SW collected the published dataset on which this analysis is based, wrote code, analysed data, prepared figures, and co-wrote the paper. GT conceived the study with SW, wrote theory, analysed data, and co-wrote the paper. Both authors gave final approval for publication and agree to be held accountable for the work performed therein.

## Acknowledgements

We thank Indira Nagesh and Inés Dawson for helpful discussions.

## Supplementary Material

### 6. List of symbols

#### Accents

^ regression estimate

– stroke average

· time derivative

#### Angles

*α* aerodynamic angle of attack

*β* angle between local velocity of wing and its surface

*γ* angle between the predicted quasi-steady aerodynamic force and the local chord

*θ* deviation angle, corresponding to elevation of wing tip in body-fixed axes

*ϕ* stroke angle, corresponding to azimuth of wing tip in body-fixed axes

*ω*_0_ wing pitch angle offset

*ω*_*r*_ linear twist gradient of wing

*ω*(*r*) local pitch angle of wing at radial coordinate *r*

#### Forces

***A*** elemental added mass force (vector)

***D***, *D* elemental quasi-steady drag force (vector, scalar)

***F*** total friction force (vector)

***L***, *L* elemental quasi-steady lift force (vector, scalar)

***P*** total pressure force (vector)

*P*_*qs*_ elemental quasi-steady pressure force (scalar)

*P*_*us*_ elemental unsteady pressure force, or added mass force (scalar)

#### Axes

*{x*_*b*_, *y*_*b*_, *z*_*b*_*}* right-handed, body-fixed axis system with origin at body centre of volume

*{x*_*L*_, *y*_*L*_, *z*_*L*_*}* right-handed, body-fixed axis system with origin at base of left wing

*{x*_*R*_, *y*_*R*_, *z*_*R*_*}* right-handed, body-fixed axis system with origin at base of right wing

*{X, Y, Z}* right-handed, lab-fixed axis system

**1**_⊥*S*_ unit vector normal to blade element surface, resolved in the body axes

**1**_‖*U*_ unit vector parallel to velocity of blade element, resolved in the body axes

**1**_⊥*U,r*_ unit vector perpendicular to blade element velocity and span, resolved in the body axes

#### Other symbols

*a*_*k*_, *b*_*k*_ coefficients of sine-cosine Fourier series

*c* wing chord length

*c*_3*/*4_, *c*_1*/*2_ distance of point back from spanwise axis (three-quarter, half-chord point)

*C*_*D*_ quasi-steady drag coefficient

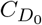 quasi-steady drag coefficient offset

*C*_*L*_ quasi-steady lift coefficient

*C*_*P*_ quasi-steady pressure force coefficient

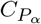 quasi-steady pressure force coefficient derivative

*f*_*n*_(*t*) harmonic function of time for *n*^th^ wingbeat

***g*** gravitational acceleration in an inertial frame of reference

*i* blade element number

*K* order of truncated Fourier series

*m* insect mass

*n* wingbeat number

*N* total number of wingbeats

*p*_0_ … *p*_3_ polynomial coefficients

*Q*_⊥*S*_ local flow speed normal to surface in an inertial frame of reference

*r* radial coordinate along wing span

***R*** rotation matrix bringing vector from the body axes into the lab axes

*R*^2^ *R*-squared statistic

*τ* discrete time

*t* continuous time

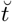 time from start of downstroke as a proportion of wingbeat period

*T*_*n*_ period of *n*^th^ wingbeat

*U* aerodynamic speed of blade element at three-quarter chord point

*U*_‖*c*_ component of velocity parallel to chord, defined at three-quarter chord point

*U*_⊥*S*_ component of velocity normal to surface, defined at three-quarter chord point

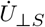 component of acceleration normal to surface at half-chord point

*V* local speed of surface

*V*_⊥*S*_ component of local velocity of wing normal to its surface

***x***_*wb*_ position vector of wing base in the body axes

***x***_3*/*4_, ***x***_1*/*2_ position of blade element in the body axes (three-quarter, half-chord point)

***X***_*b*_ position of body in the lab axes

***X***_3*/*4_, ***X***_1*/*2_ position of blade element in the lab axes (three-quarter, half-chord point)

Γ circulation around airfoil

Δ*r* blade element width

***ϵ*** error term in blade element model regression

*ρ* air density

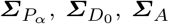 summations of blade element model predictors

## Supplementary Methods

### Kinematic transformations

We calculated the velocity of the three-quarter chord point of the *i*th blade element, 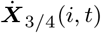, and the acceleration of the corresponding half-chord point, 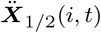, analytically. Each of these elemental quantities is defined with respect to a lab-fixed frame of reference and resolved in the lab axes, but they are obtained from the global wing and body kinematics as follows. The instantaneous position vector of the three-quarter chord point of the *i*th blade element, ***x***_3*/*4_(*i, t*), is given in the body axes by the expression:

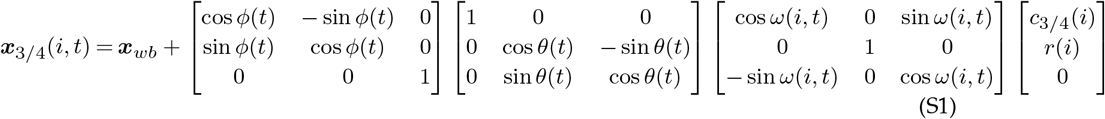

and likewise for the position vector of the corresponding half-chord point, ***x***_1*/*2_(*i, t*), substituting *c*_1*/*2_(*i*) for *c*_3*/*4_(*i*). Here, ***x***_*wb*_ is the position vector of the wing base, which we determined by finding the location of the point with the least squared variation in distance to the wing tip across all frames recorded for a given individual; *ϕ* (*t*) and *θ*(*t*) are the spherical coordinates of the wing tip in the body axes (i.e. the stroke angle and deviation angle, respectively); *ω*(*i, t*) is the instantaneous pitch angle of the *i*th blade element; *c*_3*/*4_(*i*) is the distance of the three-quarter chord point back from the spanwise axis of the wing; and *r*(*i*) is the radial coordinate of the *i*th blade element.

The position vector on the lefthand side of Eq. S1 was transformed from the body axes into the lab axes, by premultiplying it by the appropriate rotation matrix ***R***(*t*) as defined by the measured Euler angles of the body axes, then adding the position vector ***X***_*b*_ of the body axes in the lab axes:

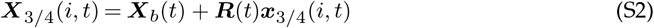

and similarly for the position vector of the corresponding half-chord point in the lab axes, ***X***_1*/*2_(*i, t*). We used Mathematica (Wolfram Mathematica, Wolfram Research Inc.) to differentiate Eqs. S1-S2 with respect to *t*, so as to derive analytical expressions for the velocity 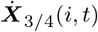 and acceleration 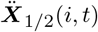. These expressions are complicated functions of the variables in Eqs. S1-S2, but are written out in full in the blade element code provided as Supporting Data S1. The net effect of these manipulations is that the velocity and acceleration of each blade element are calculated in a way that takes account of the motion both of the wings and the body. For simplicity, we drop the subscripts denoting the three-quarter chord and half-chord positions in the main text, where it is to be understood that the speed *U* (*i, t*) and acceleration *Ü* (*i, t*) of each blade element are defined at the three-quarter chord and half-chord points respectively.

We used the inverse of the rotation sequence defined by Eqs. S1-S2 to calculate the components of the blade element velocity directed parallel to the blade element chord (*U*_‖*c*_) and normal to its surface (*U*_⊥*S*_), and used these to determine the aerodynamic speed *U* (*i, t*) and angle of attack *α*(*i, t*) of each blade element according to the expressions:

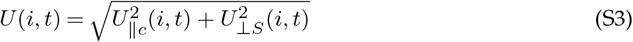

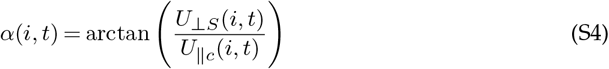

We used the same rotation sequence to determine the unit vectors **1**_⊥*U,r*_, **1**_‖*U*_, and **1**_⊥*S*_ as defined in the Methods. Matlab code implementing all of these transformations is provided as Supporting Data S1.

### Fourier series representation of the wing and body kinematics

To allow the kinematic data to be represented more compactly, we summarised the time history of each of the primary wing and body kinematic variables for each wingbeat using a harmonic function of the form:

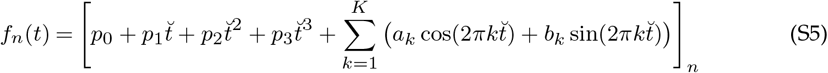

where *f*_*n*_(*t*) describes the time history of the relevant kinematic parameter for the *n*^th^ wingbeat as a function of time from the start of the downstroke (*t*), and where 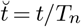denotes time as aproportion of the wingbeat period (*T*_*n*_). This expression comprises a 3^rd^ order polynomial with coefficients *p*_0_ … *p*_3_ and a *K*^th^ order Fourier series with coefficients *a*_0_ … *a*_*K*_ and *b*_0_ … *b*_*K*_. The use of a 3^rd^ order polynomial ensures that the second derivative of the function retains a linear trend term, which allows for the possibility that neither the original function nor its first and second derivatives is strictly periodic. The order of the harmonics was chosen with reference to the actual harmonic content of the data, setting *K* = 1 for the body kinematics, *K* = 4 for the wing tip kinematics, and *K* = 6 for the wing twist kinematics. Twice differentiating Eq. S5 with respect to *t*, we have:

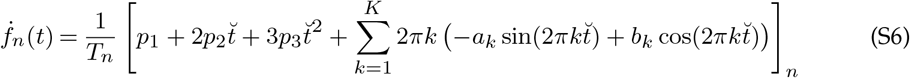

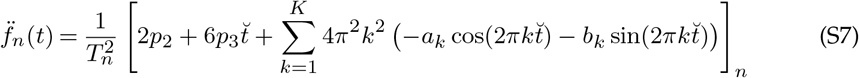

In principle, Eq. S5 can be solved in a least squares sense to determine all of the unknown coefficients. In practice, the solution is better behaved numerically across all derivatives if the higher order polynomial coefficients (*p*_2_, *p*_3_) and the Fourier series coefficients (*a*_0_ … *a*_*K*_, *b*_0_ … *b*_*K*_) are solved using Eq. S7, and then substituted in to Eq. S5 to solve for the remaining coefficients (*p*_0_, *p*_1_). Projecting the kinematic measurements into these harmonic basis functions makes a negligible difference to the numerical values that enter the blade element modelling at the discrete time points that we sampled (Fig. S1), but enables a thirty-fold compression of the data and ensures repeatability of the analysis at other resampling frequencies.

### Supplementary Video Legends

Video S1: Video showing four synchronised camera views and three-dimensional reconstruction of a hoverfly during a typical free-flight sequence. Background subtraction has been applied to improve image contrast and detail. The wings in the 3D reconstruction are coloured according to the local pitch angle *ω*(*r*); the green comet tails indicate the wing tip path. The aerodynamic forces corresponding to this particular flight sequence are plotted in Fig. 9.

Video S2: Video showing distribution of predicted resultant aerodynamic force (blue arrows) through the standard hovering wingbeat. The viewpoint rotates with the stroke, so as to look down the spanwise axis of the wing at all times. The body is shown in the background and the grey line shows the wing tip path. Individual blade elements are plotted as black lines, to show the spanwise distribution of wing twist through the wingbeat. Note that the wing is almost always highly twisted.

Video S3: Video showing distribution of predicted quasi-steady lift (blue arrows) and drag (green arrows), together with the unsteady added mass force (red arrows), through the standard hovering wingbeat. The viewpoint rotates with the stroke, so as to look down the spanwise axis of the wing at all times. The body is shown in the background and the grey line shows the wing tip path. Individual blade elements are plotted as black lines, to show the spanwise distribution of wing twist through the wingbeat. Note that the wing is almost always highly twisted.

## Supplementary Figures

**Figure S1.**
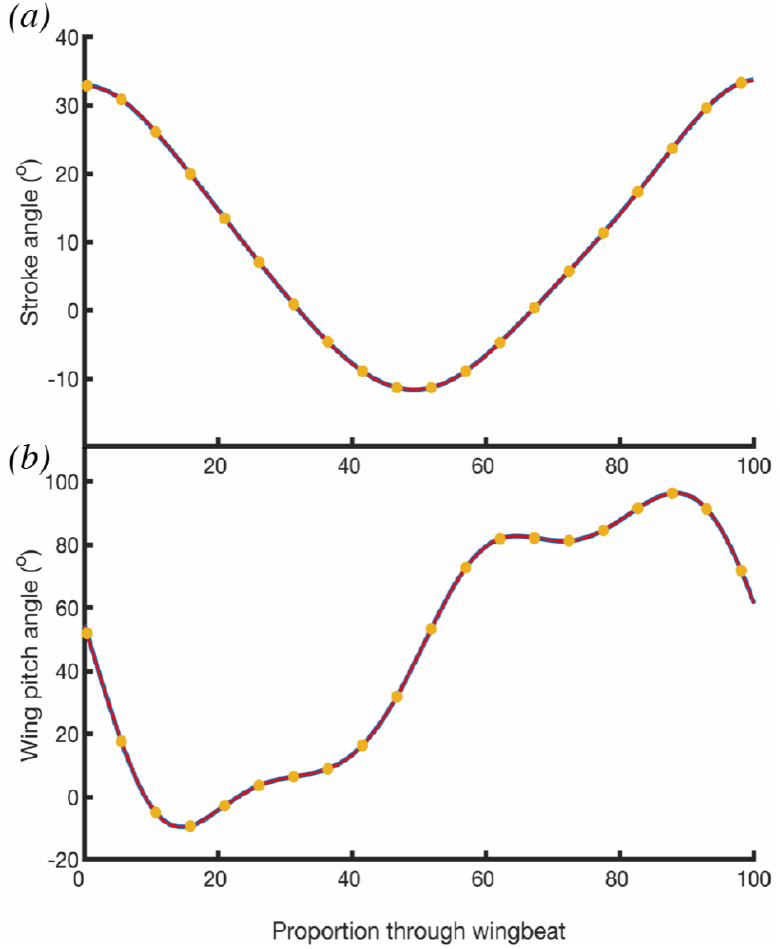
Comparison for one randomly selected wingbeat of the raw kinematic data (yellow dots), with the corresponding quintic spline fit (thick blue line), and truncated Fourier series fit (thin red line) for the stroke angle (*ϕ*) and wing pitch angle (*ω*) at mid-span. Note that these three different representations of the data that are used at successive stages of the analysis are essentially indistinguishable.

**Figure S2.**
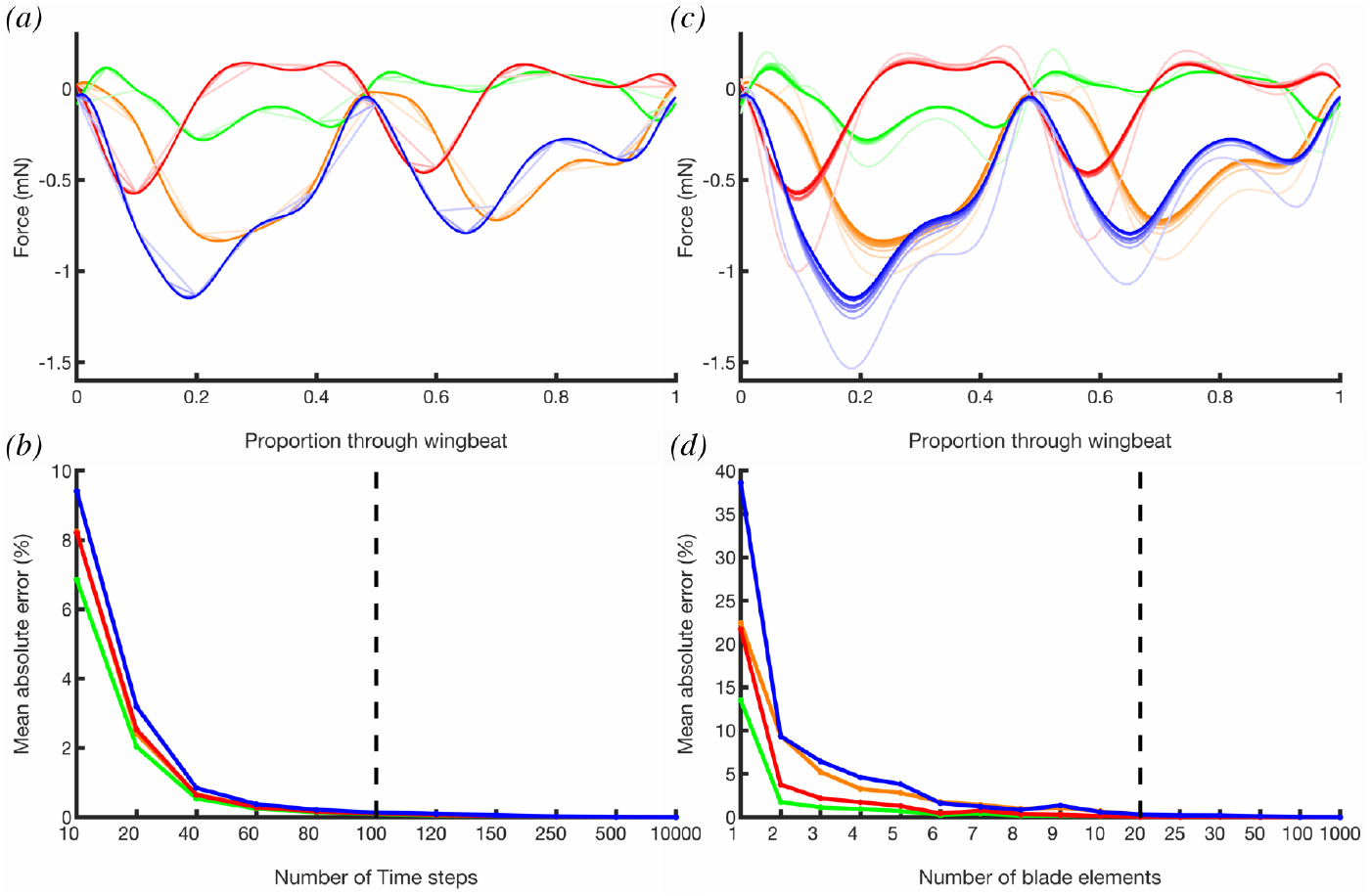
Effect of changing the number of time steps *(a-b)* and number of blade elements *(c-d)* on the aerodynamic forces predicted in the standard hovering wingbeat. *(a,c)* Predicted time-varying force components for the standard hovering wingbeat with: *(a)* 10,000 time steps and *(c)* 1,000 blade elements, superimposed with fainter lines indicating progressively reduced numbers of time steps or blade elements as shown in panels *(b,d)* below. *(b,d)* Mean absolute difference in forces compared to maxima of 10,000 time steps *(c)* or 1,000 blade elements *(d)*. Forces are shown for the right wing only, resolved in the vertical axis, and decomposed as lift (orange), drag (green), and added mass (red), together with their sum (blue). Note the nonlinear scale of the *x*-axis in *(b,d)*. Vertical black dashed lines indicate the number of time steps and blade elements that were chosen for use in the main analysis.

**Figure S3.**
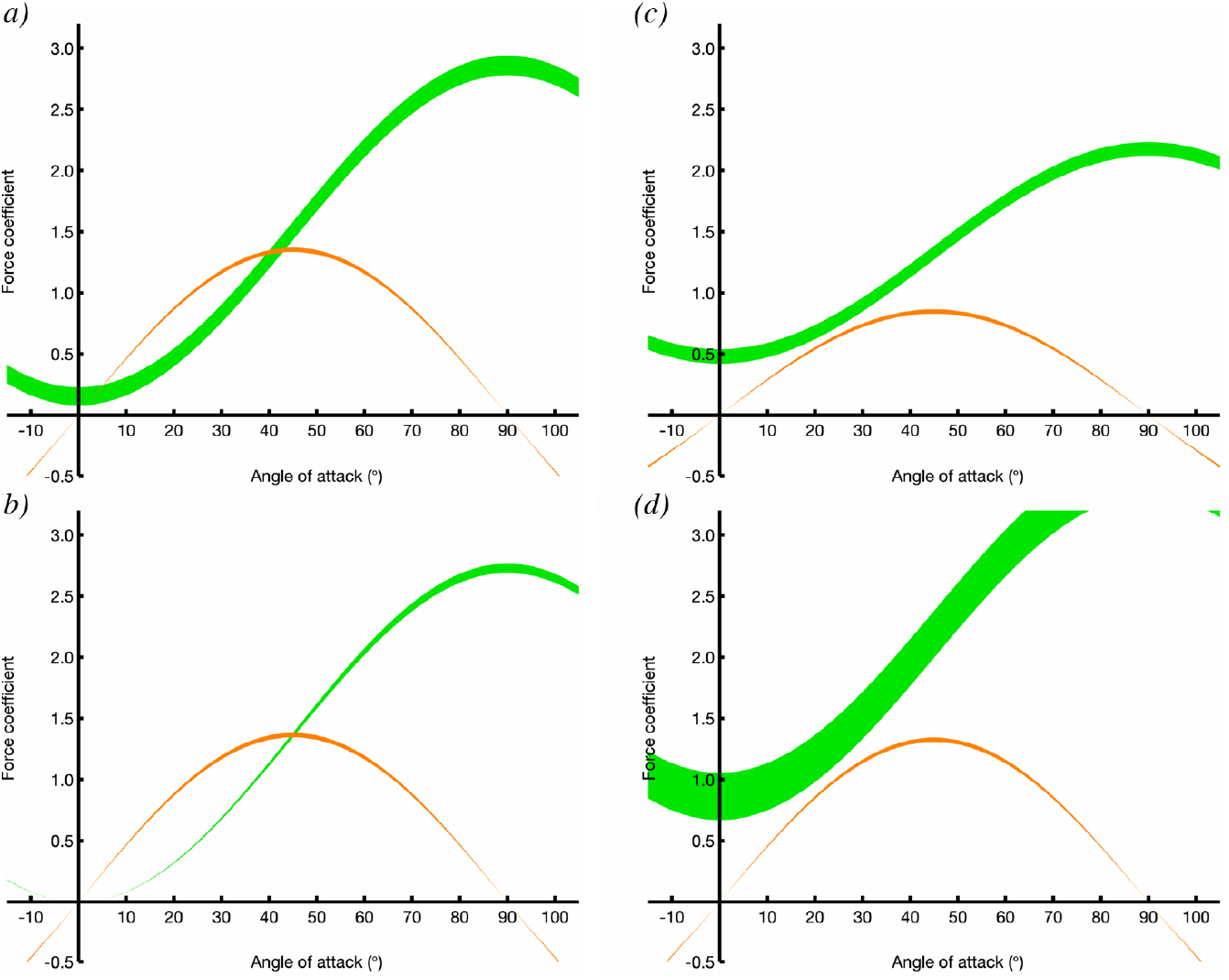
Fitted lift coefficient *Ĉ*_*L*_ (orange) and drag coefficient *Ĉ*_*D*_ (green) plotted against angle of attack *α* for the full model *(a)* and for the three simplified models described in the main text: *(b)* reduced model with no drag coefficient offset term; *(c)* wing modelled as a flat plate operating at a pitch angle matched to the measured pitch of the wing at mid-span; *(d)* model ignoring motion of the body when computing the wing kinematics. The width of the lines indicates the mean ±1SD, assessed over 10^5^ random subsamples of the data, each comprising 10% of the recorded flight sequences. Note that omitting the drag coefficient offset *(b*) makes little systematic difference to the estimated lift and drag curves, but substantially reduces the variance of the parameter estimate for the lift coefficient in random subsamples of the data, which could be beneficial if working with a smaller or noisier dataset than the one we have here. Omitting either the wing twist kinematics *(c)* or the body kinematics *(d)* from the blade element model causes a drastic change in the estimated lift and drag curves, which demonstrates the importance of accounting for these variables in the full model.

